# Dispersal rate limits range expansion rate only when it is slower than climate velocity

**DOI:** 10.64898/2026.03.16.712231

**Authors:** Nikki A. Moore, Jonathan Lenoir, Lise Comte, Jake A. Lawlor, Jennifer M. Sunday

## Abstract

Low dispersal ability might limit a species’ capacity to track its climate preferences across the landscape, yet evidence that low dispersal slows species’ range shifts under contemporary climate change remains contentious. Here we develop a new hypothesis under which we expect variation in dispersal ability to affect range expansion rates only when climate velocity *exceeds* dispersal rates, which is logical yet rarely applied because it requires a common yardstick to compare rates. We test this hypothesis using empirical relationships between dispersal ability, local range expansion rate, and the velocity of isotherm shifts in terrestrial plants and birds, all estimated in km/yr. In 370 range shifts, we found that range expansion rates were best explained by the slower of either a species’ dispersal rate or the velocity of isotherm shifts, as predicted under our hypothesis. Furthermore, when species’ dispersal rates were slower than the velocity of isotherm shifts, we found that dispersal ability positively affects range shift rates. Substantial variation in range expansion rates remained unexplained, indicating that additional factors influence range shift dynamics. Our results provide new clarity when understanding the role of dispersal ability on variation in range shift rates and emphasize the importance of evaluating dispersal capacity relative to climatic change exposure when testing hypotheses about species’ responses to ongoing environmental change.

**Significance Statement:** Species dispersal ability is widely thought to limit biodiversity redistribution in response to climate change, but we still lack a clear understanding of when or for which species dispersal limitations matter. Using hundreds of range expansion rates documented over the 10 last decades for terrestrial birds and plants together with their dispersal rates, in common units of km/yr, we show that dispersal ability slows climate-mediated range expansion, but that dispersal limitations occur only when the rates of climate change are faster than the ability of species to redistribute. So far, many species display dispersal rates higher than the velocity of climate change but dispersal limitations may become more pronounced in the future.

## Introduction

Anthropogenic climate change is expected to cause shifts in the distribution of species’ range limits across space (1). At species’ cold range edges (usually poleward and near high-elevation or low-bathymetric edges), abiotic conditions beyond species’ historic distributions are predicted to become more favourable, and species ranges are expected to locally expand as populations establish in newly-suitable areas beyond their range limits (2). Recent observations at the limits of species ranges confirm that cold, leading range edges are indeed shifting, on average, in the direction expected by climate change (3, 4). However, the extent to which observed range shifts match shifts in the distribution of isotherms across landscapes and seascapes (i.e., the velocity of climate change (5)) varies greatly (4). Anticipating species redistribution in response to climate change will require understanding why species range edges often respond idiosyncratically to changes in climate (6).

On land, species’ leading range edges tend to lag behind shifts in isotherms (7). Several factors likely contribute to these range expansion lags, such as dispersal limitation (8), limited availability (9) or accessibility (10) of suitable habitats, demographic lags (11), and differences in the biotic environment (12). The effect of dispersal in particular has received a lot of attention, likely because dispersal ability is, at least in part, a function of the measurable intrinsic traits of species (13). Yet, the extent to which dispersal limitation is responsible for variation in local range expansion rates is still unclear. Looking into the past, observations of range expansions after glacial retreat suggest that many species, even those with low estimated dispersal capacity, as per current knowledge, were able to track their climatic niches (14–16) (i.e., Reid’s Paradox (17)). However, dispersal is hypothesized to be more limiting under contemporary climate change (16) for at least two reasons: (i) current rates of climate warming are much faster than past post-glacial rates (18) and (ii) terrestrial species are currently expanding across landscapes within which habitats are far more fragmented (19). Even dispersal rates of some mammal and bird species, two highly-mobile groups, have been estimated to disperse at a slower pace than that of projected climate change (6, 20, 21). Species might be especially limited by dispersal in places where the rate of contemporary climate warming is particularly fast (like in the high arctic (22)), where the spatial temperature gradient is shallow (like at tropical latitudes (23)), or where habitat connectivity across the landscape is low.

Thus far, studies testing whether having a lower dispersal ability results in slower range expansion rates have relied on the use of proximal dispersal traits and have returned inconsistent results, both across and within taxonomic groups (13). Species’ realized dispersal rates are difficult to quantify empirically (24), vary greatly across space and time within species (25), and likely tend to underestimate the real potential for species’ dispersal, due to both the difficulty of measuring long-distance dispersal (26) and the fact that most habitats are nowadays fragmented, which renders field-based estimates of potential dispersal rates poorly reliable (19, 27). Because of this, a common approach to test for the effect of dispersal ability on range expansion across species is to use proximal traits that capture intraspecific differences in dispersal distance (such as body size (28–38), mobility (31–34), seed properties (28, 29), dispersal syndromes (39), migratory habits (28, 30, 32, 40–43), or natal dispersal distance (29)) and dispersal frequency (such as lifespan (28, 36), age at maturity (44), or the frequency of reproductive events (28, 38)), as proxies for dispersal ability. Studies that have tested whether species with traits that are expected to infer greater dispersal ability show faster range shifts (28, 30–36, 38–40, 43) or a greater tendency to shift (41, 42, 44) at cold range edges have found a combination of positive, negative, and, most often, no effect of dispersal traits on range shift rates (13).

One potential reason for the inconsistent effect of dispersal ability on observed range expansion rates might be due to the use of proximal traits. While proximal traits are useful in providing a ranking of relative dispersal ability among species considered, it is not surprising that they often fail at explaining range expansion rates. Categorical proximal traits such as migratory habit could capture variation in dispersal ability too imprecisely (45), while continuous proximal traits such as body size might scale non-linearly with realized dispersal rates (46, 47) or might correlate with other traits that also affect range dynamics but in a different direction (48). Moreover, commonly measured traits and their effect on dispersal are often not consistent across different taxonomic groups (e.g., wingspan is hypothesized to be a correlate of dispersal ability, but only applies to organisms with wings) (49), rendering it challenging to evaluate the effects of dispersal capacity on range expansion across taxa, where the bigger differences lay. Obtaining a more direct estimate of species’ potential dispersal rates rather than relying on proximal traits that differ in importance across taxa might provide new insight into the effect of dispersal ability on range expansions across species.

Another potential reason for the inconsistent relationship between proximal dispersal traits and range expansion rates is that climate change exposure is often not explicitly considered. Dispersal is expected to slow range expansion only if the rate at which a species can disperse does not allow it to keep pace with the rate at which temperature clines (i.e., isotherms) are shifting across the landscape (5, 50). If an individual of a species can disperse farther than the reach of new climatically suitable habitat beyond its range limit within its lifetime, dispersal rate would not be a limitation and should not relate to range expansion rate driven by climate change. Instead, the species’ range edge is expected to track the velocity of climate change, to the extent that other factors (such as habitat availability and accessibility) are not constraining range expansion dynamics. In other words, the effect of dispersal ability on local range expansion should depend on how fast dispersal is relative to the velocity of climate change (i.e., an estimate of the exposure to climate change in the dimension of distance per unit time). Assuming nothing else is limiting, an expanding range edge is thus theoretically expected to shift at the slower rate of either a species’ dispersal rate or the velocity of climate change, resulting in a non-linear and climate velocity-dependent relationship between species dispersal rates and their range expansion rates (Fig. 1; as has been shown in simulations (48)).

**Figure 1.**
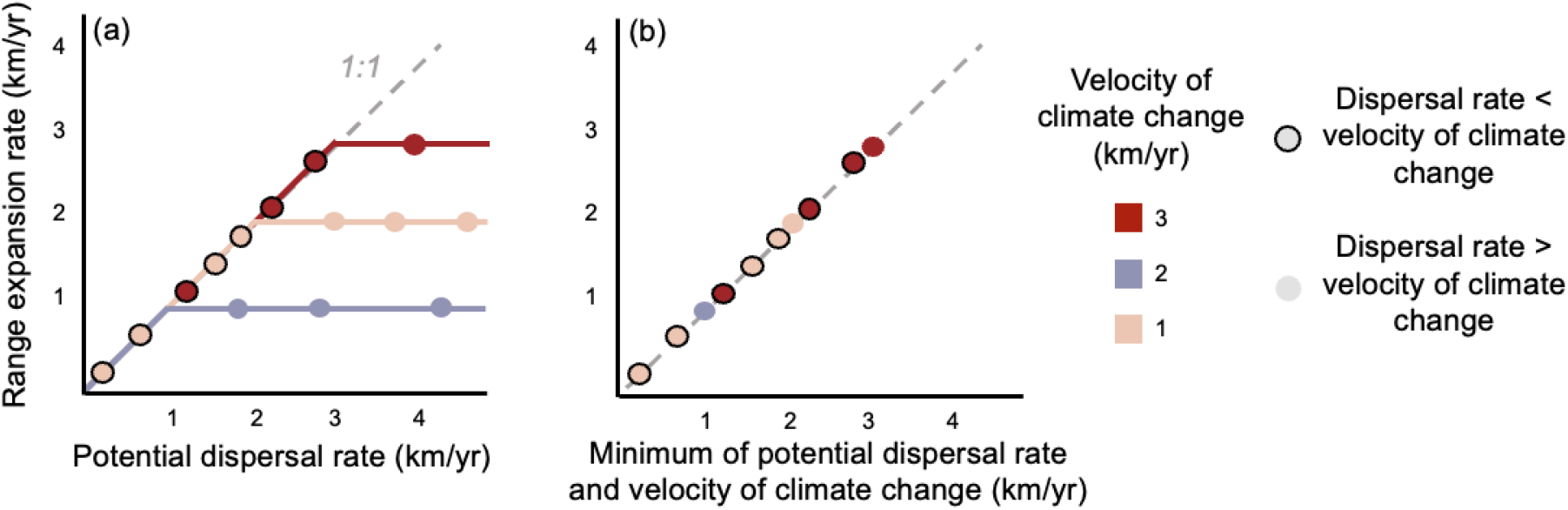
Theoretical relationship between species’ local range expansion rates, their potential dispersal rates, and the velocity of climate change. Dots represent species, while colours represent scenarios where species are exposed to different climate velocities. Dispersal ability is only expected to slow local range expansion rate if a species’ potential dispersal rate is slower than the velocity at which climatic conditions are shifting (points circled in black). When species have a potential dispersal rate that is sufficient to track this rate (non-circled points), their range is expected to expand at a rate that is lower or at best equal to the velocity of climate change rather than their dispersal rate, resulting in a non-linear relationship between potential dispersal rate and local range expansion rate (panel a). Considering the effect of the minimum rate between a species’ potential dispersal rate and the velocity of climate change on range expansion should help reveal the role of dispersal ability more clearly, since it incorporates both climate change exposure and the species’ capacity to disperse in response to climate change (panel b). Different pools of species were used in each scenario (colours) for visualization purposes, however, different relationships are expected even for the same pool of species exposed to different climate velocities.

Knowing when dispersal rate is expected to be slower than the velocity of climate change could provide improved expectations to clarify the role of dispersal ability in hindering local range expansions. If climate velocity is slower than dispersal rate (as in, for example, highly mobile taxonomic groups, such as birds shifting their ranges up mountain slopes, where climate velocity is low due to steep gradients in temperature over small spatial extents) then variation in dispersal rates should not contribute to variation in range expansion rates (blue line in Fig. 1a). However, when dispersal rate is slow compared to climate velocity (as in, for example, sessile taxonomic groups, like forest plants, shifting range edges across the lowlands, where climate velocity is high due to shallow spatial gradients in temperature), dispersal ability should be more limiting in determining range expansion rate (red line Fig. 1a). While important, this comparison is often difficult to make since proximal traits are not measured in the same units as the velocity of climate change. Indeed, without using a measure of dispersal rate that is in the same unit as the velocity of climate change (5), it is difficult to know which rate is expected to be limiting, which and is a factor that may have obscured previous investigations.

Here, for a broad set of species covering taxonomic groups with vastly different mobility, we compare empirical leading edge range shift estimates with estimates of potential dispersal rates and the velocity of isotherm shifts. Estimates of dispersal rates are based on species’ natal and propagule dispersal distances and the time before which which they disperse. We use estimate potential dispersal rates using a common yardstick (51) – units of km/yr - that can be directly compared to both estimated range shift rates and the velocity of isotherm shifts. We focus on within-generation potential dispersal rates rather than cross-generation estimates or realized movement rates across years to better capture the upper range of potential distances that a species might disperse, unconstrained by dispersal barriers. We test the hypothesis that local range expansion is slowed by dispersal ability when species cannot disperse quickly enough to keep up with the velocity at which isotherms are shifting.

## Methods

Below, we describe how we collated, filtered, and analyzed species’ estimated range shift rates at the leading edge, potential dispersal rates, and the velocity of isotherm shifts they experienced.

### Local range expansion rates

We extracted estimates of species’ range expansion rates from an updated version of the BioShifts geodatabase (data paper in prep, previous version published (52)), a global synthesis of over 30,000 geographic range shifts reported at a given range position (i.e., either the trailing/warm edge, the range centroid, or the leading/cold edge) across latitudinal or elevational gradients in both marine and terrestrial realms. Since elevational range shifts in the database are expressed as rates of vertical displacement, which would require more information about mountain topography to relate directly to horizontal dispersal rates, we extracted only data associated with latitudinal range shifts. In the database, empirical latitudinal range shift rates are reported in common units of kilometres per year (km/yr) and were assigned to a range edge (i.e., leading or trailing) based on the location of the local range edge within the study area as declared by the study. Positive range shift rates indicate that the observed range position shifted locally in the poleward direction, while negative range shift rates indicate that the observed range position shifted in the equatorward direction. The database also includes spatial polygons representing the approximate study area where each range shift estimate was reported, spatial polygons of the distribution of each species studied within that study area (see Fig. 2a), and information on the methods used to detect the range shift, including the time period during which the shift was measured.

**Figure 2.**
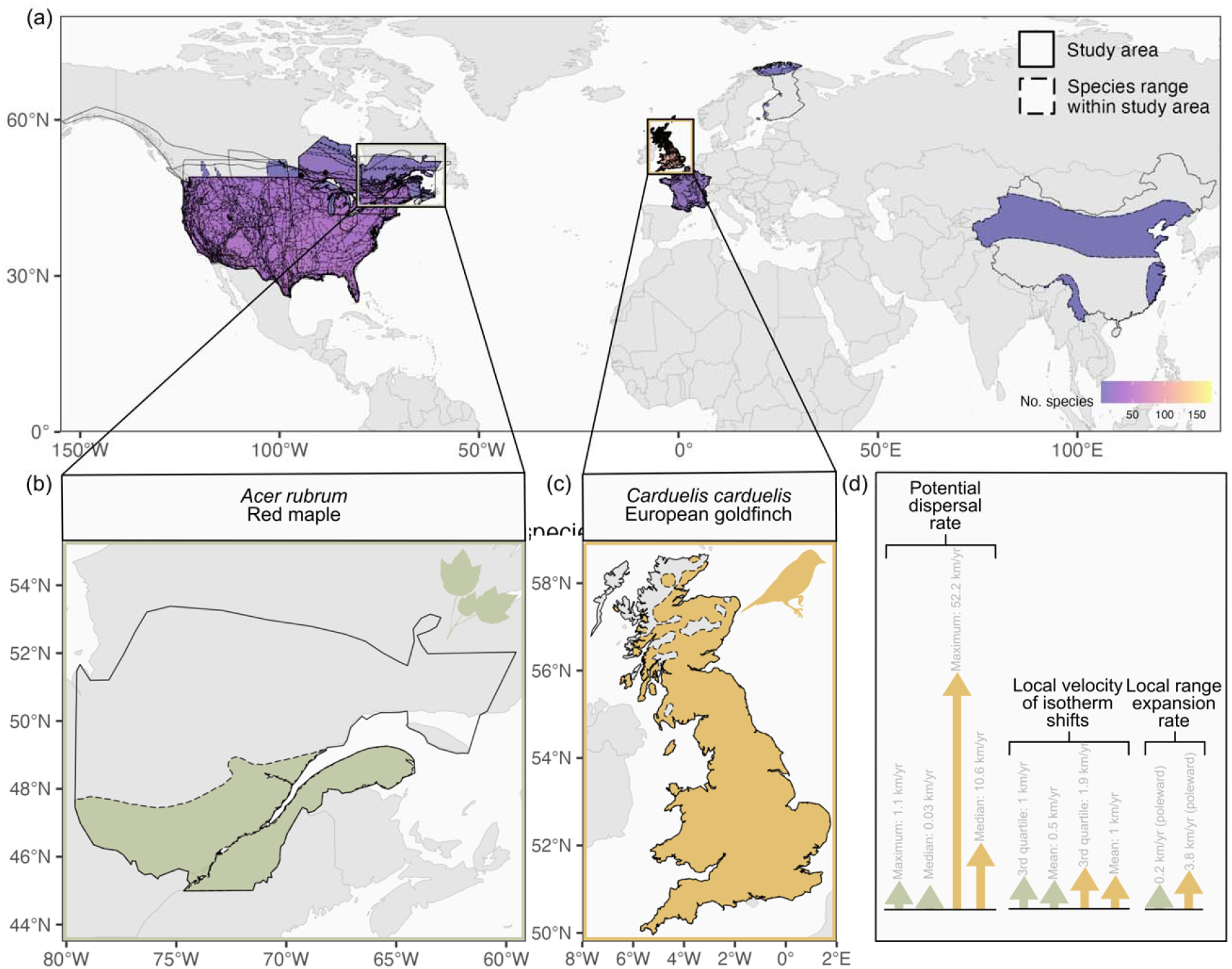
Data workflow to use a common yardstick. (a) Geographic distribution of range shift studies and (b-d) graphical representation of the data used for two exemplar species, the red maple (*Acer rubrum*) and the European goldfinch (*Carduelis carduelis*). Estimates of the leading edge (poleward) range expansion rates included in our study, which were extracted from the BioShifts geodatabase, encompass 24 study areas (polygons with solid outline in panels a-c) that were refined based on species range maps (coloured polygons with dotted outline in panels a-c). Exemplar study areas in Québec, Canada, where the local range expansion rate of the red maple was estimated (panel b, green), and in Great Britain, UK, where the local range expansion rate of the European goldfinch was estimated (panel c, orange), are shown to illustrate the data collected for both species (panel d): local range expansion rate (in km/yr from the BioShifts database), potential dispersal rate (in km/yr), and local velocity of isotherm shifts (in km/yr). We estimated species’ potential dispersal rates (maximum and median) from empirical data on species’ natal and propagule dispersal distances and the frequency at which they disperse, and we used estimates of the local velocity of isotherm shifts (3rd quartile and mean) across the portion of the study area overlapping with the species’ range (polygons with dashed outline in panels a-c).

Since our interest was in testing the role of dispersal as a limiting mechanism at expanding range fronts, we focused our study on only leading (poleward) range edge shifts where range expansion was reported. We did not include estimates of range centroid shifts since they may result from other non-dispersal related dynamics that occur throughout the species range (53) (i.e., shifts in abundance across the range, population extinction at contracting range edges), thus blurring the effect of dispersal limitation. We only considered terrestrial due to a lack of availability in comparable dispersal distance estimates for marine species.

### Potential dispersal rates

We began by collating empirical dispersal distance estimates from pre-existing databases. Because our intention was to estimate the distance that progeny have the potential to disperse in a single generation during an event that might result in population establishment, we collated databases with (i) measures of animal natal dispersal distances (54–58) (i.e., the distance between an individual’s birthplace and its site of first reproduction (59), estimated using mark-recapture methods or tracking devices) and (ii) seed or propagule dispersal distances (57, 60– 62)(i.e., the distance of seed or propagule dispersion not necessarily involving successful germination and establishment, estimated from direct observations of dispersal, dispersal of animal vectors, or dispersal experiments in a laboratory). We opted not to include estimates of dispersal distances based on short-term movements that do not usually lead to population establishment (e.g., migratory or breeding dispersal movements, foraging movements, movements within the home range (63–66)).

For each selected database, we standardized the reported taxonomy by systematically cleaning the reported scientific names and subsequently searching for their accepted scientific name and higher taxonomy from the GBIF and ITIS databases using the *traitdataform* R package (67). Once taxonomy was harmonized and standardized, we filtered each dispersal database to include only species present in the BioShifts database. We joined databases, keeping a record of the original dispersal measurement type and data source. To avoid duplicating estimates, if the accompanying publication for a dispersal database included observations from a previously published database that was in our collection, we only kept observations from the earlier published version. Finally, we retained a record of the original measurement type and data source and converted all dispersal distance measurements to common units of kilometers. Since our aim was to capture the fastest rate at which a species is capable of dispersing, and since dispersal studies often tend to underestimate dispersal (17, 68, 69), we used each species’ maximum reported dispersal distance when multiple estimates were available. However, we retained all values so that we could calculate median dispersal rates in addition to maximum to test the sensitivity of our results to using extremes.

To convert dispersal distances in kilometers to dispersal rates in km/yr, we collated data on ‘time to dispersal’ for each species by querying pre-existing databases (70–76) for estimates of the age at maturity or generation time of species for which we had dispersal distance estimates. As we did for the dispersal distance databases, we standardized the reported taxonomy before querying the databases. We chose the lowest value of generation times if multiple values were available for the same species because we were interested in maximum dispersal overall rate (and therefore earliest dispersal event). Some plant species had no available information on age at maturity or generation time but were reported as having an annual life cycle. For these species (n = 24), we inferred their dispersal frequency to be one year. We retained a record of the original measurement type and data source and converted all measurements of time to dispersal to units of years. Non-integer values were rounded up to the nearest whole year.

We defined a species’ potential dispersal rate as the speed, in km/yr, at which the progeny can successively disperse beyond the range edge. To estimate the potential dispersal rates of species, we divided estimates of dispersal distance (in kilometres) by the time between which dispersal events occur (estimated age at maturity or generation time, in years) since individuals at the expanding range front must reach maturity before dispersal of gametes or offspring can occur (17, 77). This method of calculating the potential dispersal rate is similar to approaches used by William & Blois (78) and Schloss et al. (20) to assess the extent to which dispersal is predicted to limit range shifts, except we use empirical dispersal distance observations rather than distances predicted from models that rely on proximal dispersal traits. We calculated both maximum and median potential dispersal rates based on different estimates of distance and ran models using both. Computed maximum and median potential dispersal rates, as well as the raw dispersal distances and time to dispersal estimates used to calculate them, can be found in Dataset S1.

### Local velocities of isotherm shifts

For each range expansion rate estimate, we also extracted measurements of the local velocity of isotherm shifts from the BioShifts database, which can be considered the range shift expectation based on temperature change across the species range within the focal study area. The velocities of isotherm shifts capture the rate at which local isotherms shifted during the focal study. For each range shift estimate, we therefore had estimates of isotherm shifts according to air temperatures hindcasted to the period of the study and spatially subset to the portion of the study area that overlapped with the native range of the focal species in question (referred to as ‘species-specific’ climate velocities; see Fig. 2a-c). These isotherm velocities were calculated using a gradient-based approach based on gridded mean monthly air surface temperatures (79) from the CHELSA V.1.0 and V.2.0 datasets and at different spatial resolutions (25, 50, and 110 km). This involved dividing the temporal trend in mean annual temperature in a given grid cell (i.e., the slope of a linear regression between monthly air surface temperature and time during the years where the study took place, in °C/year) by the magnitude of a vector representing the local horizontal plane in temperature (i.e., the 2-dimensional spatial gradient in temperature along latitude and longitude) across the 8 neighboring grid cells (in °C/km)(79), and extracting the latitudinal component, since latitudinal range shifts in the BioShifts database are measured as shifts along a Northward or Southward direction. We considered both the mean and a measure of a more extreme (i.e., the 3rd quartile) local velocity of isotherm shifts to which a species would be exposed across its occupied study area and repeated all analyses using both statistics.

To best capture differences in the spatial resolution at which organisms perceive and respond to environmental change, we used estimates of the velocity of isotherm shifts at the most ecologically-relevant spatial resolution for each species. The spatial resolution at which an organism can track changes in climate across their inhabited landscape likely depends on their ability to explore, sense and respond to environmental changes within that landscape throughout their lifetime (80). From the velocity of isotherm shifts calculated at a 25-km, 50-km, and 110-km spatial resolution in the BioShifts database, we assigned the most relevant spatial scale to each species according to which best matched their dispersal distance. If dispersal distance was greater than the distance between the center and any corner of the 9-grid cell squared window used to compute climate velocities at the lowest resolution (i.e., 25-km resolution grid size) we considered the next grid size up. Hence, we used the 25-km resolution grid size for species that disperse less than 47.86 km during one dispersal event (distance from center to any corner), 50-km resolution for species that disperse between 47.86 and 95.71 km, 110-km resolution for species dispersing farther than 95.71 km (see Supplementary Material Fig. S1 for distribution of isotherm velocity values at different spatial resolutions).

### Final dataset

After filtering and combining the data as described above, we retained data on potential dispersal rates and leading edge range shift rates for a total of 279 species, which were either terrestrial birds (45.52%), mammals (0.37%), plants (51.25%), amphibians (1.43%), or squamates (1.43%), and whose leading edge range shift rates were documented across 24 different study areas. Each species had at least one range shift rate estimate and sometimes more (median = 1, maximum = 5), since a range shift rate for a single species might have been estimated across several of the 24 study areas. All 478 range shift rate estimates came from scientific reports or studies covering the Northern Hemisphere solely (Fig. 2a), reflecting a commonly-observed pattern of geographical data bias in range shift detections.

Although leading (poleward) range edges are expected to shift polewards in response to climate warming at the global extent, spatial variation in climate change might result in local cooling trends or regional spatial gradients that occur in the opposite direction to the latitudinal temperature gradient, sometimes leading to the expectation of an equatorward shift at poleward range limits (i.e., expectation of a local range contraction). To ensure we included only range edge shifts expected to expand poleward, we identified and excluded range shift estimates at poleward range edges where the direction of the shift in isotherms was towards the centre of the range (i.e., poleward range edges that are expected to contract towards the centre of the range; n = 87). In addition, observations of range shifts at a given range limit do not always follow expectations and thus can occur in the opposite direction to shifts in isotherms (2). Indeed, a substantial number of range shift estimates were in the opposite direction to expectations from isotherm shifts (n = 151 negative range shift rate estimates despite positive rates of isotherm shifts). We avoided removing these range edge contractions completely, as this would constrain the error distribution (some estimation error is expected) (81), yet some contractions were so large that an ecological driver other than climate change seemed likely at play. We identified large-magnitude range edge contractions as negative shifts that were > 1 s.d. from the mean shift rate across all estimates where a poleward shift was expected (n = 21 range shift estimates, n = 16 species). We removed these from our main analysis to focus on factors that limit range expansions but provide results without this exclusion in the Supplementary Materials (Tables S4-5).

To ensure we were not removing climate change-driven range shift estimates from our main analysis, we additionally searched the literature for evidence that these extreme shifts in the opposite direction of the climate expectation were driven by non-climatic factors (findings reported in Dataset S2). Extreme range edge contractions at the leading edge were often reported for bird species, and most likely are related to habitat loss, human disturbance, or populations declining across the entire species range (e.g., the roseate tern (82), the purple martin (83), and the wood thrush (84)). Some extreme contractions were also reported for species known to have expanding ranges (e.g., the turkey vulture (85), the glossy ibis (86)) indicating dubious data points. After removing shifts where range contraction was expected (n = 87) and the few extreme cases of range edge contractions (n = 21), there did not remain sufficient sample size to test for patterns among amphibians, squamates and mammals (n = 5), so we focused our analysis on birds and plants. The geographic distribution of the assembled dataset and a graphical overview of data content can be found in Figure 2.

### Testing for relationships between range shift rates and dispersal rates

We tested the hypothesis that range shift rate responds linearly to the minimum of either a species’ potential dispersal rate or the velocity of isotherm shift. Under this hypothesis, we expected a 1:1 relationship between local range expansion rate and the minimum rate of either a species’ potential dispersal rate or the velocity of isotherm shifts (expectation 1). We also expected that the minimum rate between the two better explains variation in range expansion rates than does potential dispersal rate or the velocity of isotherm shifts alone (expectation 2); or than does their additive (expectation 3) or their interactive (expectation 4) effects. Using the rates of local range expansion as the response variable, we thus implemented 5 linear mixed-effects model types with different sets of fixed effects as predictor variables: potential dispersal rate alone, local velocity of isotherm shifts alone, minimum rate between potential dispersal rate and local velocity of isotherm shift, potential dispersal rate and local velocity of isotherm shifts together in an additive manner and next in an interactive manner. All variables were continuous measurements in units of km/yr. All models included species as a random effect term on the intercept to account for non-independence of range shift estimates made for the same species across different studies. We additionally checked for normality of residuals of all models, and for phylogenetic correlation in the residuals of the best model (see below) by calculating Pagel’s λ.

To account for uncertainty in the metrics that best capture dispersal potential (maximum or median) and the local velocity of isotherm shifts (mean or 3rd quartile), we fit the 5 model types using all possible combinations of mean and 3rd quartile of the local velocity of isotherm shifts and median and maximum potential dispersal rate, resulting in a total of 16 candidate models. We ranked all 16 candidate models by comparing model Aikake Iinformation Criterion corrected for small sample size (AICc) scores, selecting the model with the lowest AICc as the best model and considering an AICc difference greater than 2 as indicating better support (87). To assess if results were the same within each major taxonomic group (birds and plants), we refit the 16 candidate models including an additional interaction term between the taxonomic group (birds, plants) and each of the main fixed effects.

We additionally tested the hypothesis that when a species’ potential dispersal rate is slower than the local velocity of isotherm shifts, local range expansion rate is more directly influenced by species’ potential dispersal rate. Under this hypothesis, we expected there to be a linear 1:1 relationship between potential dispersal rate and local range expansion rate for observations where potential dispersal rate is slower than the local velocity of isotherm shifts (expectation 5), and for dispersal rate to explain variation in range shifts better than isotherm shift velocity (expectation 6). We tested these hypotheses by fitting 2 additional linear mixed-effects models to only estimates of local range expansion rates where species’ potential dispersal rates were equal to or slower than the local velocity of isotherm shifts, again including a random effect term of species identity on the intercept to account for non-independence between multiple observations of the same species. The first model included potential dispersal rate as a fixed effect, whereas the second model included the local velocity of isotherm shifts as a fixed effect. Models were fit with only one type of local isotherm shift velocity (maximum) and potential dispersal rate (3^rd^ quartile) as informed by model selection results of the first five model types described above.

## Results

We found that many species, and particularly birds, have potential dispersal rates that exceed the local velocity of isotherm shifts to which they were exposed (Fig. 3). After data filtration, we retained 370 range shift estimates for 224 unique species at poleward range edges (175 bird shift estimates for 91 unique species, 216 plant shift estimates for 139 unique species) for which range expansion was expected based on the direction of isotherm shift velocities (excluding extreme range contractions; see Methods). Among these range edges, species’ maximum potential dispersal rates were faster than the mean velocity of the shifting isotherms to which they were exposed 54% of the time (40% birds and 14% plants; Fig. 3 inset). This rate was higher among birds than plants. In birds, potential dispersal rate was faster than the mean velocity of isotherm shifts for 96% of bird range edges as compared to 25% of plant range edges. These comparisons were similar when comparing different parameters of each rate distribution. When the 3rd quartile of the velocity of isotherm shifts was used instead of the mean, 50% of range edges had higher dispersal compared to isotherm shift. When median, rather than maximum, potential dispersal rates were used, only a slightly lower percentage of range edges had potential dispersal rates that were faster than the velocity of isotherms shifts (49% of shifts when compared to the mean velocity [40% birds, 9% plants]; 44% shifts when compared to the 3rd quartile velocity [37% birds, 7% plants]). This suggests that for around half of all the range shifts studied, and for most bird range shifts estimated, potential dispersal rate should not be expected to limit recent range expansion rate.

**Figure 3.**
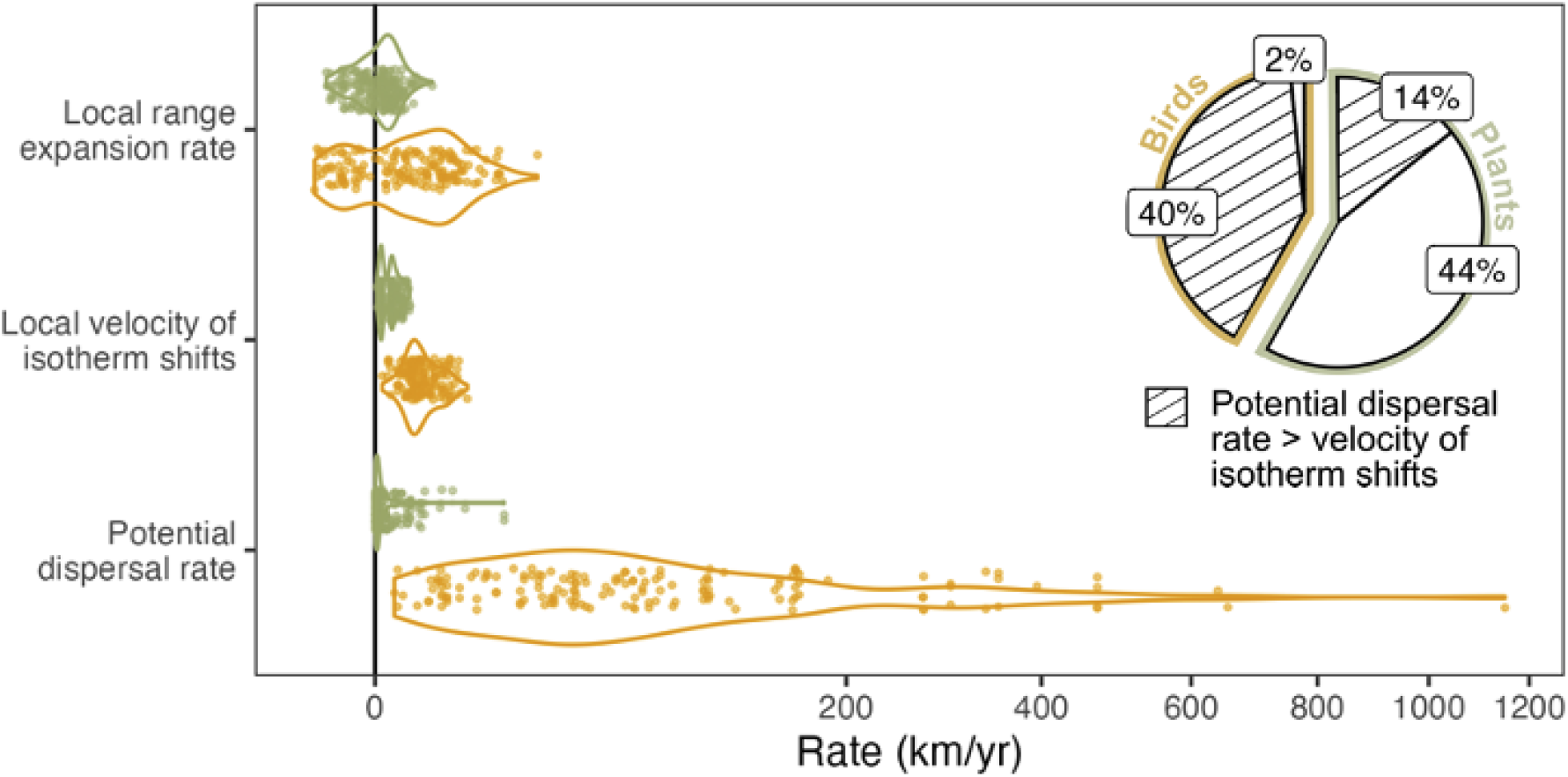
Species’ potential dispersal rates can overshoot expansion rates and climate velocities. Distributions of each rate (main panel) show that species’ maximum potential dispersal rates were often orders of magnitude higher than their estimated local range shift rates and the mean local velocity of isotherm shifts they experienced, particularly among birds (yellow) compared to plants (green). Distributions are displayed on a signed square root axis to emphasize differences in data range whilst still highlighting differences between small values. Direct comparison of species maximum potential dispersal rate to the mean local velocity of isotherm shifts show that potential dispersal rate is greater than the local velocity of isotherm shifts in 54% of the time, and mostly among bird species (inset pie chart).

We also found a poor relationship between species’ potential dispersal rates and the rates at which their leading range edge expanded (Fig. 4a, Table S1; coefficient estimate significantly deviating from 0 in only one tenth of the models). Species’ potential dispersal rates exceeded range expansion rates in half (51%) of the range shift estimates (43% when median instead of maximum potential dispersal rates was used), with potential dispersal rates sometimes being orders of magnitude greater (Fig. 3; mean ratio between magnitude of maximum potential dispersal rate and local range expansion rate = 102.43; median ratio = 1.26). While the maximum estimated range expansion rate in our study was ∼24 km/yr, 30% of species had maximum potential dispersal rates that exceeded that, with the potential to disperse at rates that ranged from 7 x 10^-5^ to 1,150 km/yr.

**Figure 4.**
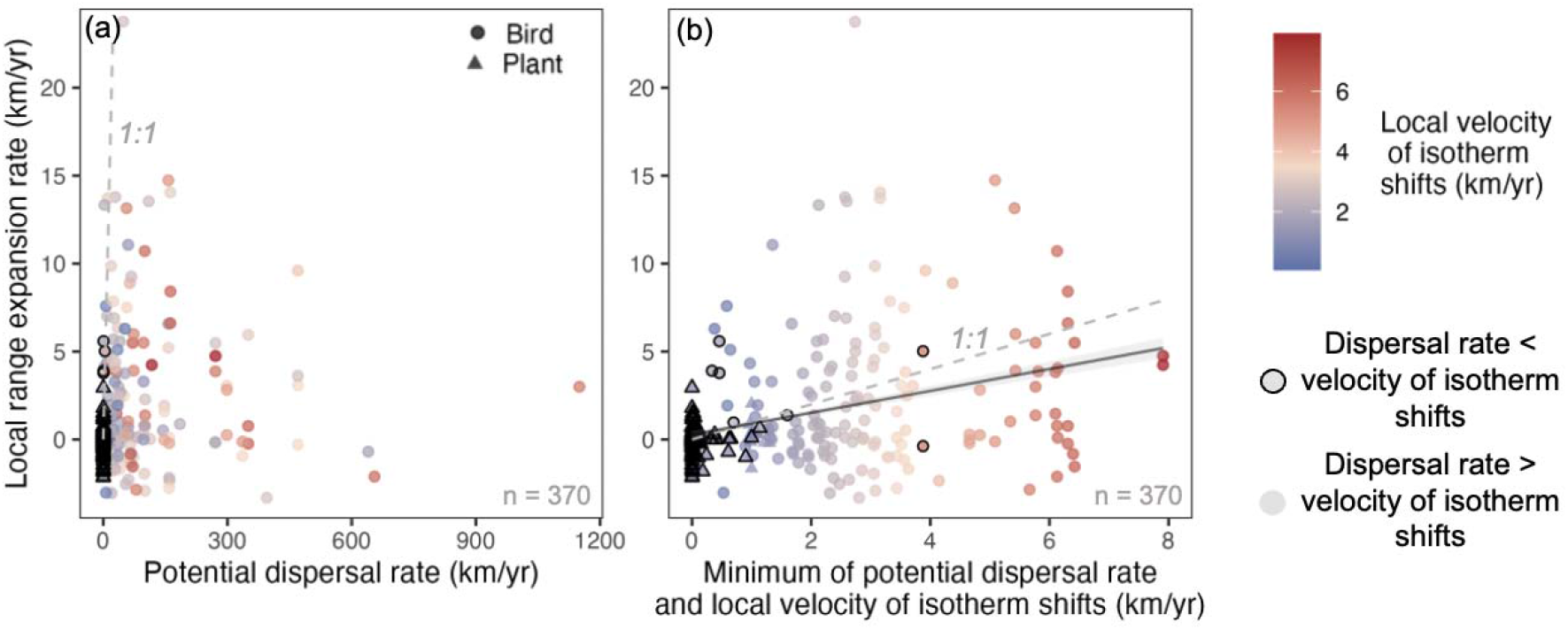
The minimum between a species’ potential dispersal rate and the local velocity of isotherm shifts explains local range expansion rate. There was no significant relationship between species’ potential dispersal rates and estimated leading edge range expansion rates (a). Instead, local range expansion rate was best explained by the minimum of a species’ potential dispersal rate and the local velocity of isotherm shifts (b). The dashed grey lines represent a 1:1 relationship, while the solid line represents predictions from the best fit linear mixed-effects model, which included the minimum of species’ maximum potential dispersal rate and the 3rd quartile of the local velocity of isotherm shifts as a fixed effect and species as a random effect term on the intercept (Table S1). Predictions represent the average effect across species, and the shaded area shows the 95% confidence interval. Points outlined in black represent local range shift estimates for which potential dispersal rate was greater than the local velocity of isotherm shift.

Range expansion rate was best explained by the minimum of a species’ potential dispersal rate and the velocity of isotherm shifts (Fig. 4b; consistent with expectation 1). The model that included the minimum rate between maximum potential dispersal rate and the 3^rd^ quartile of velocity of isotherm shifts showed better support than models that included either potential dispersal rate or the velocity of isotherm shifts alone, or the additive or interactive combinations of the two (Table S1; lowest ΔAICc = 2.13, corresponding to expectations 2-4). The minimum rate of isotherm shift velocity and potential dispersal rate had a similar estimated effect on range expansion rates as did isotherm shift velocity alone (similar parameter estimates, Table S1), but the minimum rate models had far higher cumulative weight (Table 1), and conditional R^2^ values (Table S1). Residuals of the best model were not phylogenetically correlated (i.e., λ not different from 0, phylogenetic correlation not different from white noise; Pagel’s λ = 0.01, p_0.05_ = 0.42). For models including taxonomic group (plant or bird) as an interactive term in the model’s formula, the minimum rate candidate model remained within the top models set but no clear best model could be identified (Table S2).

**Table 1.** Summary of model fits across different model types.

| Model type | Minimum model rank | Mean model rank | Cumulative weight |
| --- | --- | --- | --- |
| Minimum rate | 1 | 5.5 | 0.91 |
| Potential dispersal rate and local velocity of isotherm shifts (interactive) | 3 | 7.25 | 0.04 |
| Local velocity of isotherm shifts | 4 | 8 | 0.02 |
| Potential dispersal rate and local velocity of isotherm shifts (additive) | 5 | 9.5 | 0.02 |
| Potential dispersal rate | 15 | 15.5 | 0.00 |

Note: Model ranks and weights based on AICc score of 16 individual models are summarized within each of the five main model types. Models within model type varied in specific parameters used (i.e., the 3rd quartile versus mean local velocity of isotherm shift, median versus maximum potential dispersal rate; see Table S1 for individual model results).

When we considered only range shift estimates for which the maximum potential dispersal rate was slower than the velocity of isotherm shifts (n = 185), the range expansion rate was better explained by potential dispersal rate than the velocity of isotherm shifts. We found a positive relationship between range expansion rate and maximum potential dispersal rate with a slope coefficient of 0.63±0.11 (Fig. 5a, Table 2; using 3rd quartile of velocity of isotherm shift). While range expansion rate also showed a positive relationship with the velocity of isotherm shifts for this subset of range shift estimates (Fig. 5b, Table 2), the slope was lower (slope coefficient of 0.28±0.09, corresponding to expectation 5). In addition, the model including potential dispersal rate alone had a lower AICc score and a higher explanatory power (ΔAICc = 5.36, higher R^2^ in Table 2, corresponding to expectation 6).

**Table 2.**
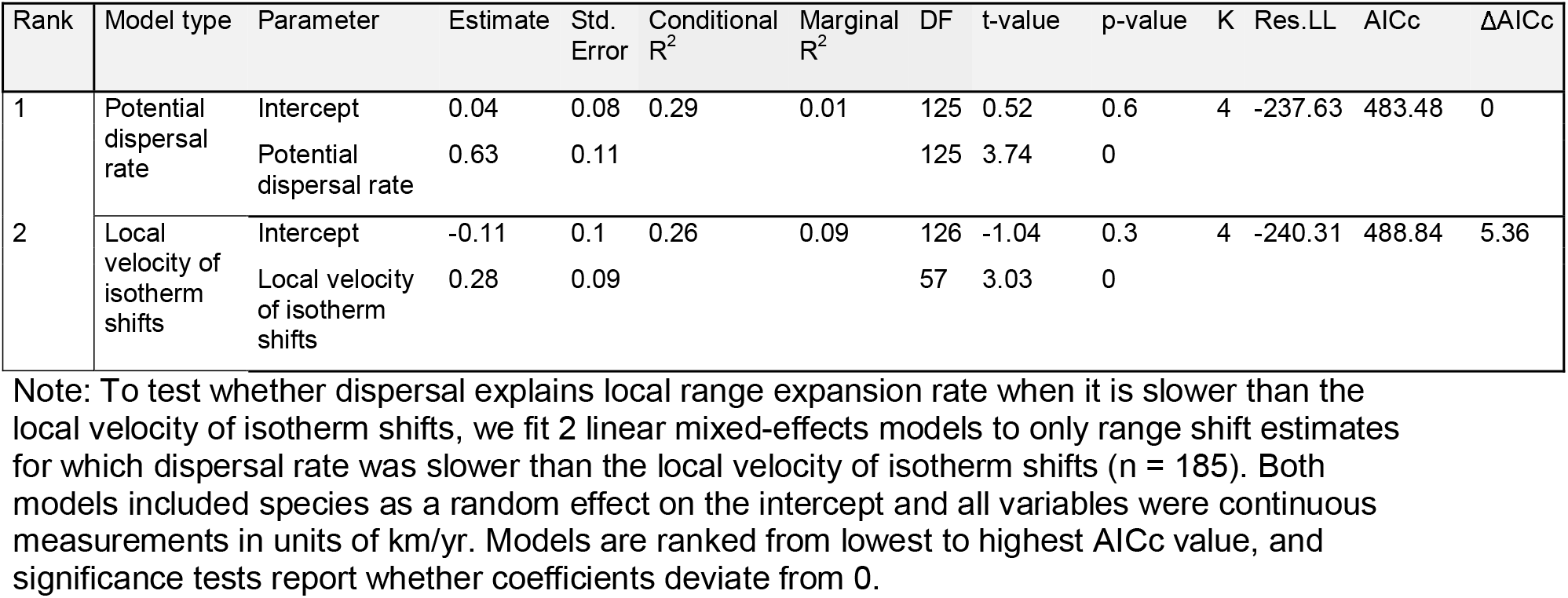
Summary of linear mixed-effects models fit to local range expansion rates for which potential dispersal rate was slower than the local velocity of isotherm shifts.

**Figure 5.**
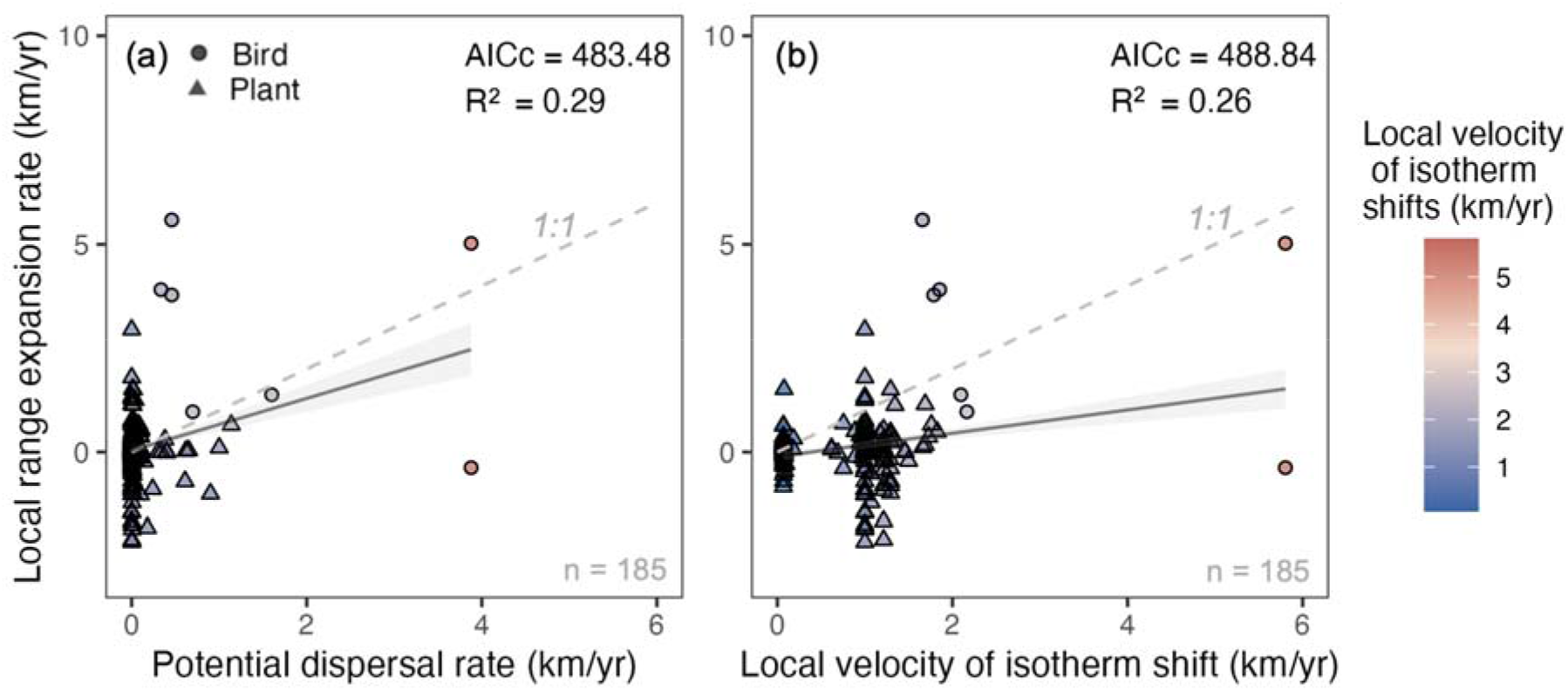
Local range expansion rate was best explained by species’ potential dispersal rates when species’ potential dispersal rates were slower than the local velocity of isotherm shifts. Among range shift estimates where species’ potential dispersal rate was slower than the local velocity of isotherm shifts, local range expansion rate increased with species’ potential dispersal rate (a). Although the relationship between local range expansion rate and the local velocity of isotherm shifts was similar (b), the model including potential dispersal rate had a lower AICc score, a higher conditional R^2^ value, and had a slope coefficient closer to the 1:1 expected relationship (dotted line). Solid lines represent predictions from linear mixed-effects models fit to local range expansion rate, including either potential dispersal rate (a) or the local velocity of isotherm shifts (b) (Table S1) as a fixed effect and species as a random effect on the intercept. Shaded areas show the 95% confidence interval. Model outputs shown here use measurements of the 3rd quartile of the local velocity of isotherm shifts and species’ maximum potential dispersal rates.

Including the few estimates of extreme local range edge contraction (n = 21) at the leading range edge led to poorly fit models (i.e., highest marginal R^2^ values of 0.05; Table S4) and less clear results (i.e., up to seven candidate models competing as the best model [ΔAICc < 2] and effect size were generally smaller; Table S4). Although the minimum rate model was no longer the only top model and was not the model with the lowest overall and mean rank, it appeared in the top model set (i.e., models with ΔAICc < 2; Table S4).

## Discussion

Our results show that species locally expanded their ranges according to their potential dispersal abilities, but that dispersal limitations occurred only when the velocity of isotherm shifts was higher than the potential ability of species to redistribute. Through comparing potential dispersal rates directly with velocities of isotherm shifts and species’ range expansion rates in the same units, we provide clear evidence that dispersal limitation can contribute to range expansion lags, but that many species display dispersal abilities currently sufficient to keep up with the shifting isotherms. By using trait values that are more direct estimates of the process of dispersal, setting out clear theoretical expectations (48), and considering climate change exposure when testing hypotheses (13, 28), our ‘common yardstick’ approach brings needed clarity to the role of dispersal limitation in slowing species’ range shifts. Below, we discuss our findings and how they can advance the science of understanding and predicting biodiversity redistribution in response to climate change.

The weak direct effect of potential dispersal rates on local range expansion rates when considering all observations in our study can be explained by the fact that many species in our dataset, including most birds and some plants, had potential dispersal rates that were orders of magnitude faster than the velocities of isotherm shifts. By converting potential dispersal abilities to units of velocity, it was apparent even before fitting any models that potential dispersal rate was not likely to explain variation in range expansion rates, since most species could disperse at rates well above that which was required to track changes in their thermal preferences across the area in which their leading range edge shift was estimated. This large difference in magnitude between potential dispersal rates and velocities of isotherm shifts can help explain why the minimum of the two rates were consistently better at explaining range expansions than dispersal or velocity of isotherm shifts alone; the minimum rate brought some range shift expectations down from enormously high dispersal rates (e.g., 270 km/year for *Anas platyrhynchos*, the mallard) to values that match isotherm shift velocity (e.g., 3.1 km/year for that same species). Overall, these results emphasize the importance of considering potential dispersal rate relative to local climate change exposure when predicting species’ responses, and that improving our understanding of range-shift trait relationships requires testing for non-linear or complex relationships (88).

Our comparison of maximum dispersal rate to range expansion rates also underscores the idea that *potential* dispersal rates do not necessarily equate to *realized* dispersal rates. Many species with sufficiently high dispersal abilities to theoretically track isotherm shifts still lagged behind the 1:1 expectation between isotherm shift rates and realized range expansion rates (non-outlined points under 1:1 line in Fig. 4b, for which climate velocity is essentially the x-axis). This highlights the difference between realized dispersal rates as observed during range expansion over a few generations (study durations range from 12 to 96 years), and potential dispersal rates, which were estimated here by combining empirical observations of dispersal distances with reproductive frequencies. This difference could be explained by stochasticity in establishment success, variation in landscape features that affect dispersal (e.g., wind), habitat fragmentation or low availability of suitable habitats (19), or inaccuracies of dispersal rates that were not collected from the same population or region which we had estimates of range expansion rates. Over longer time periods it is possible that realized range expansion rates could ‘catch up’ to better match climate velocities during more conducive conditions to allow maximum dispersal, even if those conditions are short or infrequently met. A similar mechanism was part of the explanation for Reid’s Paradox (17). When the speed of range expansion during the Holocene was faster than expected among many tree species based on predictions according to dispersal rates across shorter time scales (akin to “realized range” shifts in this study), this could be explained by occurrence of long-distance dispersal events that manifested only over longer time frames. A large review of range shift studies also found that range shifts estimated across longer study durations were better matched with predictions from climate velocities (2). Realized dispersal is not simply diffusive population spread at a steady rate but is expected to be dynamic due to differing ecology, varying conditions, and random events.

Our findings highlight the importance of considering potential dispersal relative to climate change exposure and help to make sense of inconsistencies across previous studies that tested for an effect of dispersal on range expansion rates. Our findings would predict that studies testing for a relationship between dispersal traits and range expansion rates among highly mobile and dispersive taxa, which are more likely to have dispersal rates that surpass velocities of climate change, are less likely to find an effect than those testing for a relationship among less dispersive taxa. While the use of different traits and methods of measuring range shifts at a given range position makes it difficult to compare results among previous studies, the expectation that dispersal traits are more likely to influence range expansion among generally less dispersive species seems to be supported in the literature. Of the studies reviewed in MacLean and Beissinger (13) that tested the relationship between proxy of dispersal traits and terrestrial latitudinal range expansion at the leading range edge, a positive effect of dispersal traits on range expansion was found more often among taxa who have more limited mobility or rely on passive dispersal mechanisms (insects and plants; 2/3 (34, 37, 42)), while a negative, inconsistent, or no relationship was often found among birds (7/10 (28–30, 30, 33, 40, 42, 43, 89, 90)). For example, birds with greater mean natal dispersal distance showed a reduced probability of expanding their latitudinal range in Britain (29), while butterflies with large wingspan and high mobility showed greater range expansions in Finland (34). Extending our results to range expansions across elevational gradients, we can expect the velocity of temperature change across elevational gradients to be substantially lower (i.e., lower distances required to track isotherm shifts per unit time) than across latitudinal gradients, due to steep spatial gradients in temperature (5). Here, even fewer taxa are expected to be dispersal-limited (i.e., to have dispersal rates lower than isotherm shift velocities). Indeed, most studies reviewed in MacLean and Beissinger found an unexpected or inconsistent effect of dispersal traits on elevational range shifts at the leading edge of terrestrial species, even among plants (5/6 (28, 39–41, 91)).

While our results agreed with theoretical expectations, our models left substantial unexplained variation. Our best model explained ∼ 41% of variation in local range expansion rates, with only ∼14% being attributed to the minimum rate between potential dispersal rate and the velocity of isotherm shifts. Although our theoretical expectations were based on a scenario where either isotherm shift velocity or dispersal rate limits range expansion, we expected deviations from this simple expectation due to other factors involved in range dynamics (e.g., habitat availability (9), low recruitment or survival beyond the range edge (11), range limits uncoupled from thermal niche limits (92), responses to changes in other niche dimensions (41)), the stochastic nature of dispersal (24), and complex interactions between dispersal ability and habitat connectivity across the landscape (19, 93). Additionally, some unexplained variation among range expansion rates is undoubtedly due to estimation error in all three rate estimates. Estimating the velocity of climate change is imperfect (94) and sensitive to methodology (95), as is the detection and quantification of range edge shifts (2) and dispersal potential (24). The lack of a clear best model when relationships were allowed to vary between birds and plants is also likely related to differences in expectations between groups; it is unsurprising that the minimum rate model did not outcompete models with only potential dispersal rate or the velocity of isotherm shift when group was included as an interactive term in the models since the minimum rate was almost always dispersal rate for plants and the rate of isotherm shifts for birds. To detect a better fit of a model with conditional effects of dispersal rate and isotherm shifts within groups, there must be sufficient variation in dispersal rate such that the minimum rate is sometimes dispersal rate and sometimes the rate of isotherm shift. In the context of species redistribution, going from theory to predictive power using trait-based approaches still represents a daunting task, but this study provides a proof of concept that there may be a path forward.

Interestingly, extremes were more important than means when explaining range shift rate variation. The best minimum rate model was composed of rates derived from a species’ maximum potential dispersal rate or the upper 3rd quartile of the velocity of isotherm shifts within the study area. The better performance of maximum versus average dispersal capacity agrees with previous findings of a higher-than-expected speed of post-glacial range shifts that could be explained by way of extreme, long-distance events rather than diffusion based on mean dispersal rates at an expanding population front (i.e., a long-tailed dispersal kernel; Reid’s Paradox (26)). Our estimates were not derived from dispersal kernels based on probabilities, however they do incorporate population variation in dispersal distances, and so capture some relevant variation in species’ dispersal capacity. This might be especially necessary given that empirical dispersal distance estimates tend to give underestimates in both means and extremes (96). In the calculation of the velocity of isotherm shifts, statistics for each species were derived from the variation across grid cells of the species range where the study was conducted, with some study domains being very large. Our findings suggest that using a maximum estimate of climate change exposure was better than using a mean, possibly because the mean averages across grid cells when the relevant range edge shifts could be taking place in specific locations where more extreme changes are occurring within the study area. Indeed, extreme weather and climate events can trigger range expansion or even act to amplify it by changing the dispersal patterns of a species, increasing the probability and frequency of long-distance dispersal events (97). While previous syntheses have shown that range shifts often do not match the direction or magnitude of the mean climate expectation across study areas (2), our results suggest it is worth considering whether the magnitude and direction of shifts fall within the range of possible expectations across the space where the range shift was observed.

Testing hypotheses in ecology often requires converting information about processes that are studied separately into the same ecological currencies. Here, we converted dispersal ability, temperature change exposure, and range edge shift observations into the same units so that they could be directly compared, allowing more concrete expectations and new insights into whether dispersal is expected to be limiting. This revealed that dispersal ability across plants and birds is clearly highly variable, and in many cases (mostly in birds) has been more than sufficient to track shifts in temperature across the landscape. Although it requires extra effort and access to species-level traits data, our results demonstrate the value of estimating traits that more directly capture the mechanisms of range shifts rather than ones that are proximal to the process in question, thus improving the potential of traits to help understand variation in species range shifts (48). With these unified currencies, range expansions were best predicted by the minimum between potential dispersal rate and the velocity of temperature change in the models we tested. In some cases, remaining lags in species’ expansion rates relative to the velocity of temperature change could diminish over time as species “catch up” to isotherms given more time to express their potential dispersal rates, while in other cases they might increase as the rate of climate warming continues to accelerate (98) and as habitat fragmentation and loss affect the realized dispersal rate. Continued observations of the dynamics of species’ ranges set against clear expectations based on a common yardstick and on knowledge on natural history and habitat availability will be key to disentangling these futures.

## Acknowledgments

N.A.M. is funded by a Doctoral Canada National Research Scholarship from the Natural Sciences and Engineering Research Council of Canada. This research was a product of the BIOSHIFTS working group, funded by the Centre for the Synthesis and Analysis of Biodiversity (CESAB), a key program of the French Foundation for Research on Biodiversity (FRB; www.fondationbiodiversite.fr). We are grateful for all BIOSHIFT members’ support.

