## Supporting Information for "Dispersal rate limits range expansion rate only when it is slower than climate velocity"

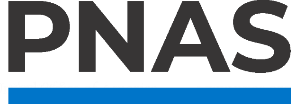

**Supporting Information for**

Dispersal rate limits range expansions only when it is slower than the velocity of climate change

Nikki A. Moore, Jonathan Lenoir, Lise Comte, Jennifer M. Sunday

Nikki A. Moore

**This PDF file includes:**

Figure

Tables S1 to S5

**Other supporting materials for this manuscript include the following:**

Datasets S1 to S2

Figures

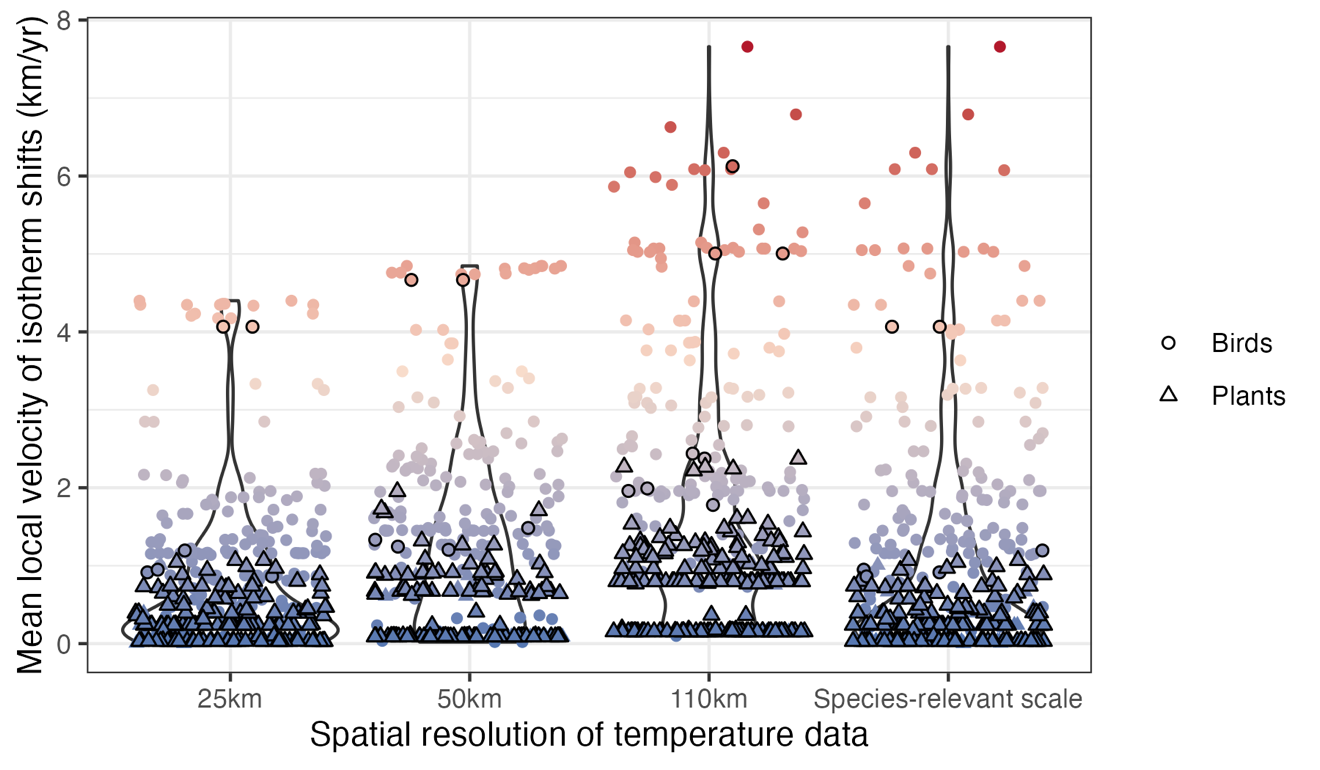

Fig. S1. The local velocity of isotherm shifts depends on the spatial resolution of temperature data. The magnitude of isotherm shift velocity tends to be greater when the spatial resolution of temperature data used to calculate the velocity is more coarse (larger spatial resolution).

Tables

Table S1. Summary of linear mixed effect models fit to local range expansion rates

| RankRank | Model type | Velocity of isotherm shifts | Dispersal distance | Parameter | Estimate | Std.Error | Conditional R^2^ | Marignal R^2^ | DF | t-value | p-value | K | LL | AICc | | ΔAICc | AIC weight |
| --- | --- | --- | --- | --- | --- | --- | --- | --- | --- | --- | --- | --- | --- | --- | --- | --- | --- |
| 1 | Minimum rate | 3rd quartile | Max | Intercept | 0.28 | 0.2 | 0.27 | 0.14 | 223 | 1.41 | 0.16 | 4 | -913.43 | | 1834.96 | 0 | 0.68 |
|  |  |  |  | Minimum rate | 0.62 | 0.09 |  |  | 145 | 7.16 | 0 |  |  | |  |  |  |
| 2 | Minimum rate | 3rd quartile | Median | Intercept | 0.32 | 0.2 | 0.27 | 0.13 | 223 | 1.66 | 0.1 | 4 | -914.49 | | 1837.1 | 2.13 | 0.23 |
|  |  |  |  | Minimum rate | 0.61 | 0.09 |  |  | 145 | 6.96 | 0 |  |  | |  |  |  |
| 3 | Potential dispersal rate: interactive | 3rd quartile | Max | Intercept | -0.08 | 0.24 | 0.28 | 0.13 | 222 | -0.34 | 0.73 | 6 | -914.69 | | 1841.6 | 6.64 | 0.02 |
|  |  |  |  | Potential dispersal rate | 0.01 | 0 |  |  | 222 | 1.33 | 0.19 |  |  | |  |  |  |
|  |  |  |  | Local velocity of isotherm shifs | 0.73 | 0.11 |  |  | 144 | 6.45 | 0 |  |  | |  |  |  |
|  |  |  |  | Potential dispersal rate: Local velocity of isotherm shifts | 0 | 0 |  |  | 144 | -1.94 | 0.05 |  |  | |  |  |  |
| 4 | Velocity of isotherm shift | 3rd quartile | - | Intercept | 0.06 | 0.23 | 0.26 | 0.11 | 223 | 0.26 | 0.79 | 4 | -917.05 | | 1842.2 | 7.24 | 0.02 |
|  |  |  |  | Local velocity of isotherm shifts | 0.62 | 0.1 |  |  | 145 | 6.53 | 0 |  |  | |  |  |  |
| 5 | Potential dispersal rate: additive | 3rd quartile | Median | Intercept | 0.06 | 0.23 | 0.27 | 0.11 | 222 | 0.27 | 0.79 | 5 | -916.53 | | 1843.23 | 8.27 | 0.01 |
|  |  |  |  | Potential dispersal rate | 0 | 0 |  |  | 222 | -1.02 | 0.31 |  |  | |  |  |  |
|  |  |  |  | Local velocity of isotherm shifts | 0.65 | 0.1 |  |  | 145 | 6.48 | 0 |  |  | |  |  |  |
| 6 | Potential dispersal rate: additive | 3rd quartile | Max | Intercept | 0.04 | 0.23 | 0.26 | 0.11 | 222 | 0.19 | 0.85 | 5 | -916.58 | | 1843.33 | 8.37 | 0.01 |
|  |  |  |  | Potential dispersal rate | 0 | 0 |  |  | 222 | -0.96 | 0.34 |  |  | |  |  |  |
|  |  |  |  | Local velocity of isotherm shifts | 0.67 | 0.11 |  |  | 145 | 6.14 | 0 |  |  | |  |  |  |
| 7 | Potential dispersal rate: interactive | 3rd quartile | Median | Intercept | -0.01 | 0.23 | 0.27 | 0.12 | 222 | -0.04 | 0.97 | 6 | -915.62 | | 1843.46 | 8.5 | 0.01 |
|  |  |  |  | Potential dispersal rate | 0 | 0.01 |  |  | 222 | 0.82 | 0.42 |  |  | |  |  |  |
|  |  |  |  | Local velocity of isotherm shifts | 0.71 | 0.11 |  |  | 144 | 6.54 | 0 |  |  | |  |  |  |
|  |  |  |  | Potential dispersal rate: Local velocity of isotherm shifts | 0 | 0 |  |  | 144 | -1.34 | 0.18 |  |  | |  |  |  |
| 8 | Potential dispersal rate: interactive | Mean | Max | Intercept | 0.2 | 0.21 | 0.29 | 0.12 | 222 | 0.93 | 0.35 | 6 | -915.92 | | 1844.08 | 9.12 | 0.01 |
|  |  |  |  | Potential dispersal rate | 0.01 | 0 |  |  | 222 | 2.24 | 0.03 |  |  | |  |  |  |
|  |  |  |  | Local velocity of isotherm shifts | 0.93 | 0.15 |  |  | 144 | 6.19 | 0 |  |  | |  |  |  |
|  |  |  |  | Potential dispersal rate: Local velocity of isotherm shifts | 0 | 0 |  |  | 144 | -3.06 | 0 |  |  | |  |  |  |
| 9 | Minimum rate | Mean | Median | Intercept | 0.49 | 0.19 | 0.26 | 0.1 | 223 | 2.56 | 0.01 | 4 | -919.48 | | 1847.08 | 12.12 | 0 |
|  |  |  |  | Minimum rate | 0.67 | 0.11 |  |  | 145 | 6.05 | 0 |  |  | |  |  |  |
| 10 | Minimum rate | Mean | Max | Intercept | 0.48 | 0.19 | 0.26 | 0.1 | 223 | 2.49 | 0.01 | 4 | -919.51 | | 1847.13 | 12.17 | 0 |
|  |  |  |  | Minimum rate | 0.67 | 0.11 |  |  | 145 | 6.04 | 0 |  |  | |  |  |  |
| 11 | Potential dispersal rate: interactive | Mean | Median | Intercept | 0.32 | 0.21 | 0.28 | 0.11 | 222 | 1.51 | 0.13 | 6 | -918.35 | | 1848.93 | 13.97 | 0 |
|  |  |  |  | Potential dispersal rate | 0 | 0 |  |  | 222 | 1 | 0.32 |  |  | |  |  |  |
|  |  |  |  | Local velocity of isotherm shifts | 0.84 | 0.14 |  |  | 144 | 6 | 0 |  |  | |  |  |  |
|  |  |  |  | Potential dispersal rate: Local velocity of isotherm shifts | 0 | 0 |  |  | 144 | -1.96 | 0.05 |  |  | |  |  |  |
| 12 | Local velocity of isotherm shift | Mean | - | Intercept | 0.41 | 0.2 | 0.26 | 0.09 | 223 | 1.99 | 0.05 | 4 | -920.85 | | 1849.81 | 14.85 | 0 |
|  |  |  |  | Local velocity of isotherm shifts | 0.66 | 0.11 |  |  | 145 | 5.77 | 0 |  |  | |  |  |  |
| 13 | Potential dispersal rate: additive | Mean | Median | Intercept | 0.42 | 0.2 | 0.27 | 0.09 | 222 | 2.03 | 0.04 | 5 | -920.29 | | 1850.74 | 15.78 | 0 |
|  |  |  |  | Potential dispersal rate | 0 | 0 |  |  | 222 | -1.06 | 0.29 |  |  | |  |  |  |
|  |  |  |  | Local velocity of isotherm shifts | 0.71 | 0.12 |  |  | 145 | 5.75 | 0 |  |  | |  |  |  |
| 14 | Potential dispersal rate: additive | Mean | Max | Intercept | 0.41 | 0.2 | 0.27 | 0.09 | 222 | 2 | 0.05 | 5 | -920.6 | | 1851.37 | 16.41 | 0 |
|  |  |  |  | Potential dispersal rate | 0 | 0 | x | 0.09 | 222 | -0.7 | 0.48 |  |  | |  |  |  |
|  |  |  |  | Local velocity of isotherm shifts | 0.71 | 0.13 | 0.27 | 0.09 | 145 | 5.33 | 0 |  |  | |  |  |  |
| 15 | Potential dispersal rate | - | Max | Intercept | 0.94 | 0.19 | 0.29 | 0.01 | 222 | 4.94 | 0 | 4 | -933.98 | | 1876.08 | 41.11 | 0 |
|  |  |  |  | Potential dispersal rate | 0 | 0 |  |  | 222 | 2 | 0.05 |  |  | |  |  |  |
| 16 | Potential dispersal rate | - | Median | Intercept | 1.03 | 0.19 | 0.3 | 0 | 222 | 5.42 | 0 | 4 | -935.58 | | 1879.26 | 44.3 | 0 |
|  |  |  |  | Potential dispersal rate | 0 | 0 |  |  | 222 | 0.84 | 0.4 |  |  | |  |  |  |

Note: A total of 16 linear mixed effect models of 5 different model types were fit to species’ estimated range shift rates at poleward range edges where local range expansion was expected based on the direction of isotherm velocity, after excluding extreme negative shifts (n = 370; n = 21 extreme negative shifts removed). The 5 model types included the following fixed effects: potential dispersal rate alone, local velocity of isotherm shifts alone, the minimum rate between potential dispersal rate and velocity of isotherm shift, potential dispersal rate and local velocity of isotherm shifts together in an additive manner, and potential dispersal rate and local velocity of isotherm shifts together in an interactive manner. All models included species as a random effect on the intercept and all variables were continuous measurements in units of km/yr. Models are ranked from lowest to highest AICc value, and significance tests report whether coefficients deviate from 0.

Table S2. Summary of linear mixed effect models fit to local range expansion rates including an interaction with taxonomic group

| Rank | Model type | Velocity of isotherm shifts | Dispersal rate | Parameter | Estimate | Std.Error | Conditional R^2^ | Marginal R^2^ | DF | t-value | p-value | K | LL | AICc | ΔAICc | AIC weight |
| --- | --- | --- | --- | --- | --- | --- | --- | --- | --- | --- | --- | --- | --- | --- | --- | --- |
| 1 | Potential dispersal rate (additive) | 3rd quartile | Median | Intercept | 2.24 | 0.49 | 0.32 | 0.18 | 220 | 4.62 | 0 | 7 | -904.05 | 1822.42 | 0 | 0.16 |
|  |  |  |  | Potential dispersal rate | 0 | 0 |  |  | 220 | -2.01 | 0.05 |  |  |  |  |  |
|  |  |  |  | Group: plants | -2.36 | 0.47 |  |  | 220 | -5.04 | 0 |  |  |  |  |  |
|  |  |  |  | Local velocity of isotherm shifts | 0.21 | 0.13 |  |  | 145 | 1.56 | 0.12 |  |  |  |  |  |
|  |  |  |  | Potential dispersal rate:group (plants) | 0.02 | 0.13 |  |  | 220 | 0.18 | 0.85 |  |  |  |  |  |
| 2 | Potential dispersal rate | - | Median | Intercept | 2.87 | 0.28 | 0.32 | 0.18 | 220 | 10.28 | 0 | 6 | -905.3 | 1822.83 | 0.41 | 0.13 |
|  |  |  |  | Potential dispersal rate | 0 | 0 |  |  | 220 | -1.86 | 0.06 |  |  |  |  |  |
|  |  |  |  | Group: plants | -2.86 | 0.35 |  |  | 220 | -8.17 | 0 |  |  |  |  |  |
|  |  |  |  | Potential dispersal rate:group (plants) | 0.02 | 0.13 |  |  | 220 | 0.17 | 0.86 |  |  |  |  |  |
| 3 | Potential dispersal rate (additive) | Mean | Median | Intercept | 2.47 | 0.41 | 0.32 | 0.18 | 220 | 6.03 | 0 | 7 | -904.37 | 1823.05 | 0.63 | 0.12 |
|  |  |  |  | Potential dispersal rate | -0.01 | 0 |  |  | 220 | -2.08 | 0.04 |  |  |  |  |  |
|  |  |  |  | Group: plants | -2.5 | 0.44 |  |  | 220 | -5.72 | 0 |  |  |  |  |  |
|  |  |  |  | Local velocity of isotherm shifts | 0.2 | 0.15 |  |  | 145 | 1.35 | 0.18 |  |  |  |  |  |
|  |  |  |  | Potential dispersal rate:group (plants) | 0.03 | 0.13 |  |  | 220 | 0.19 | 0.85 |  |  |  |  |  |
| 4 | Potential dispersal rate (additive) | 3rd quartile | Max | Intercept | 2.16 | 0.48 | 0.31 | 0.18 | 220 | 4.48 | 0 | 7 | -904.5 | 1823.31 | 0.89 | 0.1 |
|  |  |  |  | Potential dispersal rate | 0 | 0 |  |  | 220 | -1.76 | 0.08 |  |  |  |  |  |
|  |  |  |  | Group: plants | -2.31 | 0.47 |  |  | 220 | -4.95 | 0 |  |  |  |  |  |
|  |  |  |  | Local velocity of isotherm shifts | 0.25 | 0.14 |  |  | 145 | 1.81 | 0.07 |  |  |  |  |  |
|  |  |  |  | Potential dispersal rate:group (plants) | 0.03 | 0.13 |  |  | 220 | 0.27 | 0.79 |  |  |  |  |  |
| 5 | Local velocity of isotherm shifts | 3rd quartile | - | Intercept | 1.95 | 0.5 | 0.3 | 0.18 | 222 | 3.86 | 0 | 6 | -905.76 | 1823.75 | 1.33 | 0.08 |
|  |  |  |  | Local velocity of isotherm shifts | 0.22 | 0.14 |  |  | 144 | 1.56 | 0.12 |  |  |  |  |  |
|  |  |  |  | Group: plants | -1.88 | 0.59 |  |  | 222 | -3.17 | 0 |  |  |  |  |  |
|  |  |  |  | Local velocity of isotherm shifts:group (plants) | -0.3 | 0.39 |  |  | 144 | -0.76 | 0.45 |  |  |  |  |  |
| 6 | Minimum rate | 3rd quartile | Max | Intercept | 2 | 0.49 | 0.3 | 0.18 | 222 | 4.06 | 0 | 6 | -905.88 | 1823.98 | 1.56 | 0.07 |
|  |  |  |  | Minimum rate | 0.21 | 0.14 |  |  | 144 | 1.5 | 0.14 |  |  |  |  |  |
|  |  |  |  | Group: plants | -1.98 | 0.54 |  |  | 222 | -3.67 | 0 |  |  |  |  |  |
|  |  |  |  | Minimum rate:group (plants) | -0.21 | 0.8 |  |  | 144 | -0.26 | 0.79 |  |  |  |  |  |
| 7 | Potential dispersal rate (additive) | Mean | Max | Intercept | 2.43 | 0.41 | 0.32 | 0.18 | 220 | 5.95 | 0 | 7 | -905.03 | 1824.38 | 1.96 | 0.06 |
|  |  |  |  | Potential dispersal rate | 0 | 0 |  |  |  | -1.71 | 0.09 |  |  |  |  |  |
|  |  |  |  | Group: plants | -2.48 | 0.44 |  |  |  | -5.65 | 0 |  |  |  |  |  |
|  |  |  |  | Local velocity of isotherm shifts | 0.23 | 0.15 |  |  |  | 1.49 | 0.14 |  |  |  |  |  |
|  |  |  |  | Potential dispersal rate:group (plants) | 0.03 | 0.13 |  |  |  | 0.26 | 0.79 |  |  |  |  |  |
| 8 | Potential dispersal rate | - | Max | Intercept | 2.85 | 0.3 | 0.31 | 0.18 | 220 | 9.57 | 0 | 6 | -906.15 | 1824.54 | 2.12 | 0.06 |
|  |  |  |  | Potential dispersal rate | 0 | 0 |  |  | 220 | -1.31 | 0.19 |  |  |  |  |  |
|  |  |  |  | Group: plants | -2.84 | 0.37 |  |  | 220 | -7.76 | 0 |  |  |  |  |  |
|  |  |  |  | Potential dispersal rate:group (plants) | 0.03 | 0.13 |  |  | 220 | 0.24 | 0.81 |  |  |  |  |  |
| 9 | Minimum rate | 3rd quartile | Median | Intercept | 2.1 | 0.48 | 0.3 | 0.18 | 222 | 4.4 | 0 | 6 | -906.17 | 1824.57 | 2.16 | 0.06 |
|  |  |  |  | Minimum rate | 0.18 | 0.14 |  |  | 144 | 1.28 | 0.2 |  |  |  |  |  |
|  |  |  |  | Group: plants | -2.09 | 0.52 |  |  | 222 | -3.98 | 0 |  |  |  |  |  |
|  |  |  |  | Minimum rate:group (plants) | -0.18 | 1.17 |  |  | 144 | -0.15 | 0.88 |  |  |  |  |  |
| 10 | Local velocity of isotherm shifts | Mean | - | Intercept | 2.31 | 0.41 | 0.3 | 0.17 | 222 | 5.67 | 0 | 6 | -906.49 | 1825.21 | 2.8 | 0.04 |
|  |  |  |  | Local velocity of isotherm shifts | 0.15 | 0.15 |  |  | 144 | 1.01 | 0.32 | 6 |  |  |  |  |
|  |  |  |  | Group: plants | -2.3 | 0.49 |  |  | 222 | -4.66 | 0 | 6 |  |  |  |  |
|  |  |  |  | Local velocity of isotherm shifts:group (plants) | -0.1 | 0.8 |  |  | 144 | -0.13 | 0.9 | 6 |  |  |  |  |
| 11 | Minimum rate | Mean | Median | Intercept | 2.32 | 0.4 | 0.3 | 0.17 | 222 | 5.79 | 0 | 6 | -906.52 | 1825.26 | 2.84 | 0.04 |
|  |  |  |  | Minimum rate | 0.14 | 0.15 |  |  | 144 | 0.98 | 0.33 |  |  |  |  |  |
|  |  |  |  | Group: plants | -2.29 | 0.46 |  |  | 222 | -4.99 | 0 |  |  |  |  |  |
|  |  |  |  | Minimum rate:group (plants) | -0.53 | 3.03 |  |  | 144 | -0.17 | 0.86 |  |  |  |  |  |
| 12 | Minimum rate | Mean | Max | Intercept | 2.32 | 0.4 | 0.3 | 0.17 | 222 | 5.75 | 0 | 6 | -906.52 | 1825.28 | 2.86 | 0.04 |
|  |  |  |  | Minimum rate | 0.14 | 0.15 |  |  | 144 | 0.98 | 0.33 |  |  |  |  |  |
|  |  |  |  | Group: plants | -2.31 | 0.46 |  |  | 222 | -5.01 | 0 |  |  |  |  |  |
|  |  |  |  | Minimum rate:group (plants) | -0.07 | 1.75 |  |  | 144 | -0.04 | 0.97 |  |  |  |  |  |
| 13 | Potential dispersal rate (interactive) | 3rd quartile | Median | Intercept | 2.05 | 0.55 | 0.32 | 0.19 | 220 | 3.75 | 0 | 10 | -903.64 | 1827.89 | 5.47 | 0.01 |
|  |  |  |  | Potential dispersal rate | 0 | 0.01 |  |  | 220 | -0.51 | 0.61 |  |  |  |  |  |
|  |  |  |  | Local velocity of isotherm shifts | 0.27 | 0.16 |  |  | 142 | 1.7 | 0.09 |  |  |  |  |  |
|  |  |  |  | Group: plants | -1.99 | 0.63 |  |  | 220 | -3.14 | 0 |  |  |  |  |  |
|  |  |  |  | Potential dispersal rate: Local of isotherm shifts | 0 | 0 |  |  | 142 | -0.31 | 0.76 |  |  |  |  |  |
|  |  |  |  | Potential dispersal rate:group (plants) | 0.03 | 0.19 |  |  | 220 | 0.17 | 0.86 |  |  |  |  |  |
|  |  |  |  | Local velocity of isotherm shifts:group (plants) | -0.35 | 0.4 |  |  | 142 | -0.87 | 0.39 |  |  |  |  |  |
|  |  |  |  | Potential dispersal rate: Local of isotherm shifts:group (plants) | -0.02 | 0.21 |  |  | 142 | -0.08 | 0.94 |  |  |  |  |  |
| 14 | Potential dispersal rate (interactive) | 3rd quartile | Max | Intercept | 2.03 | 0.61 | 0.31 | 0.18 | 220 | 3.35 | 0 | 10 | -904.02 | 1828.65 | 6.23 | 0.01 |
|  |  |  |  | Potential dispersal rate | 0 | 0 |  |  | 220 | -0.77 | 0.44 |  |  |  |  |  |
|  |  |  |  | Local velocity of isotherm shifts | 0.29 | 0.18 |  |  | 142 | 1.68 | 0.09 |  |  |  |  |  |
|  |  |  |  | Group: plants | -1.98 | 0.69 |  |  | 220 | -2.88 | 0 |  |  |  |  |  |
|  |  |  |  | Potential dispersal rate: Local of isotherm shifts | 0 | 0 |  |  | 142 | 0.07 | 0.95 |  |  |  |  |  |
|  |  |  |  | Potential dispersal rate:group (plants) | 0.05 | 0.18 |  |  | 220 | 0.27 | 0.79 |  |  |  |  |  |
|  |  |  |  | Local velocity of isotherm shifts:group (plants) | -0.36 | 0.42 |  |  | 142 | -0.87 | 0.38 |  |  |  |  |  |
|  |  |  |  | Potential dispersal rate: Local of isotherm shifts:group (plants) | -0.03 | 0.2 |  |  | 142 | -0.13 | 0.89 |  |  |  |  |  |
| 15 | Potential dispersal rate (interactive) | Mean | Median | Intercept | 2.41 | 0.45 | 0.32 | 0.18 | 220 | 5.33 | 0 | 10 | -904.31 | 1829.23 | 6.81 | 0.01 |
|  |  |  |  | Potential dispersal rate | 0 | 0 |  |  | 220 | -0.9 | 0.37 |  |  |  |  |  |
|  |  |  |  | Local velocity of isotherm shifts | 0.23 | 0.18 |  |  | 142 | 1.25 | 0.21 |  |  |  |  |  |
|  |  |  |  | Group: plants | -2.41 | 0.54 |  |  | 220 | -4.5 | 0 |  |  |  |  |  |
|  |  |  |  | Potential dispersal rate: Local of isotherm shifts | 0 | 0 |  |  | 142 | -0.24 | 0.81 |  |  |  |  |  |
|  |  |  |  | Potential dispersal rate:group (plants) | 0.04 | 0.19 |  |  | 220 | 0.23 | 0.82 |  |  |  |  |  |
|  |  |  |  | Local velocity of isotherm shifts:group (plants) | -0.18 | 0.82 |  |  | 142 | -0.21 | 0.83 |  |  |  |  |  |
|  |  |  |  | Potential dispersal rate: Local of isotherm shifts:group (plants) | -0.1 | 0.66 |  |  | 142 | -0.15 | 0.88 |  |  |  |  |  |
| 16 | Potential dispersal rate (interactive) | Mean | Max | Intercept | 2.29 | 0.51 | 0.32 | 0.18 | 220 | 4.51 | 0 | 10 | -904.9 | 1830.41 | 7.99 | 0 |
|  |  |  |  | Potential dispersal rate | 0 | 0 |  |  | 220 | -0.39 | 0.7 |  |  |  |  |  |
|  |  |  |  | Local velocity of isotherm shifts | 0.29 | 0.21 |  |  | 142 | 1.41 | 0.16 |  |  |  |  |  |
|  |  |  |  | Group: plants | -2.3 | 0.59 |  |  | 220 | -3.93 | 0 |  |  |  |  |  |
|  |  |  |  | Potential dispersal rate: Local of isotherm shifts | 0 | 0 |  |  | 142 | -0.43 | 0.67 |  |  |  |  |  |
|  |  |  |  | Potential dispersal rate:group (plants) | 0.05 | 0.18 |  |  | 220 | 0.29 | 0.77 |  |  |  |  |  |
|  |  |  |  | Local velocity of isotherm shifts:group (plants) | -0.22 | 0.84 |  |  | 142 | -0.26 | 0.79 |  |  |  |  |  |
|  |  |  |  | Potential dispersal rate: Local of isotherm shifts:group (plants) | -0.11 | 0.62 |  |  | 142 | -0.17 | 0.86 |  |  |  |  |  |

Note: To assess if results were the same within each major taxonomic group (birds and plants), we refit the 16 candidate models of 5 model types to species’ estimated local range expansion rates with an additional interaction between group (birds, plants) and each of the main fixed effects (n = 370). All models included species as a random effect on the intercept and all variables were continuous measurements in units of km/yr. Models are ranked from lowest to highest AICc value, and significance tests report whether coefficients deviate from 0.

Table S3. Summary of model fits across different model types for the set of models including group as an interaction term

| Model type | Minimum model rank | Mean model rank | Cumulative weight |
| --- | --- | --- | --- |
| Potential dispersal rate and local velocity of isotherm shifts (additive) | 1 | 3.75 | 0.44 |
| Potential dispersal rate | 2 | 5.0 | 0.19 |
| Local velocity of isotherm shifts | 5 | 7.5 | 0.12 |
| Minimum rate | 6 | 9.5 | 0.21 |
| Potential dispersal rate and local velocity of isotherm shifts (interactive) | 13 | 14.5 | 0.03 |

Note: Model ranks and weights based on AIC score of 16 individual models are summarized within each of the five main model types. Models within model type varied in specific parameters used (i.e., 3rd quantile versus mean velocity of isotherm shift, median versus maximum potential dispersal rate; see Table S2 for individual model results).

**Table S4.** Summary of linear mixed effect models fit to local range expansion rates including extreme contractions

| Rank | Model type | Velocity of isotherm shifts | Dispersal rate | Parameter | Estimate | Std.Error | Conditional R^2^ | Marginal R^2^ | DF | t-value | p-value | K | LL | AICc | ΔAICc | AIC weight |
| --- | --- | --- | --- | --- | --- | --- | --- | --- | --- | --- | --- | --- | --- | --- | --- | --- |
| 1 | Potential dispersal rate (additive) | Mean | Median | Intercept | 0.05 | 0.27 | 0.26 | 0.05 | 228 | 0.18 | 0.86 | 5 | -1084.12 | 2178.39 | 0 | 0.2 |
|  |  |  |  | Potential dispersal rate | -0.01 | 0 |  |  | 228 | -1.9 | 0.06 |  |  |  |  |  |
|  |  |  |  | Local velocity of isotherm shifts | 0.66 | 0.16 |  |  | 160 | 4.09 | 0 |  |  |  |  |  |
| 2 | Potential dispersal rate (additive) | Mean | Maximum | Intercept | 0.03 | 0.27 | 0.25 | 0.04 | 228 | 0.13 | 0.9 | 5 | -1084.55 | 2179.27 | 0.88 | 0.13 |
|  |  |  |  | Potential dispersal rate | 0 | 0 | 0.25 | 0.04 | 228 | -1.65 | 0.1 |  |  |  |  |  |
|  |  |  |  | Local velocity of isotherm shifts | 0.7 | 0.18 | 0.25 | 0.04 | 160 | 3.98 | 0 |  |  |  |  |  |
| 3 | Potential dispersal rate (interactive) | Mean | Maximum | Intercept | -0.09 | 0.29 | 0.25 | 0.05 | 228 | -0.32 | 0.75 | 6 | -1083.67 | 2179.55 | 1.16 | 0.11 |
|  |  |  |  | Potential dispersal rate | 0 | 0 |  |  | 228 | 0.27 | 0.79 |  |  |  |  |  |
|  |  |  |  | Local velocity of isotherm shifts | 0.82 | 0.2 |  |  | 159 | 4.14 | 0 |  |  |  |  |  |
|  |  |  |  | Potential dispersal rate:local velocity of isotherm shifts | 0 | 0 |  |  | 159 | -1.33 | 0.19 |  |  |  |  |  |
| 4 | Potential dispersal rate (interactive) | Mean | Median | Intercept | -0.01 | 0.28 | 0.25 | 0.05 | 228 | -0.04 | 0.97 | 6 | -1083.75 | 2179.71 | 1.33 | 0.1 |
|  |  |  |  | Potential dispersal rate | 0 | 0.01 | 0.25 | 0.05 | 228 | -0.41 | 0.68 |  |  |  |  |  |
|  |  |  |  | Local velocity of isotherm shifts | 0.74 | 0.19 | 0.25 | 0.05 | 159 | 3.98 | 0 |  |  |  |  |  |
|  |  |  |  | Potential dispersal rate:local velocity of isotherm shifts | 0 | 0 | 0.25 | 0.05 | 159 | -0.85 | 0.4 |  |  |  |  |  |
| 5 | Minimum rate | Mean | Maximum | Intercept | 0.1 | 0.26 | 0.24 | 0.04 | 229 | 0.4 | 0.69 | 4 | -1085.81 | 2179.72 | 1.33 | 0.1 |
|  |  |  |  | Minimum rate | 0.53 | 0.15 |  |  | 160 | 3.65 | 0 |  |  |  |  |  |
| 6 | Minimum rate | Mean | Median | Intercept | 0.11 | 0.26 | 0.24 | 0.04 | 229 | 0.44 | 0.66 | 4 | -1085.88 | 2179.87 | 1.48 | 0.1 |
|  |  |  |  | Minimum rate | 0.53 | 0.15 | 0.24 | 0.04 | 160 | 3.63 | 0 |  |  |  |  |  |
| 7 | Local velocity of isotherm shifts | Mean | - | Intercept | 0.03 | 0.27 | 0.24 | 0.04 | 229 | 0.1 | 0.92 | 4 | -1085.91 | 2179.93 | 1.54 | 0.09 |
|  |  |  |  | Local velocity of isotherm shifts | 0.55 | 0.15 | 0.24 | 0.04 | 160 | 3.62 | 0 |  |  |  |  |  |
| 8 | Minimum rate | 3rd quartile | Maximum | Intercept | 0.07 | 0.27 | 0.21 | 0.03 | 229 | 0.26 | 0.79 | 4 | -1086.89 | 2181.89 | 3.5 | 0.04 |
|  |  |  |  | Minimum rate | 0.39 | 0.12 | 0.21 | 0.03 | 160 | 3.31 | 0 |  |  |  |  |  |
| 9 | Potential dispersal rate (additive) | 3rd quartile | Median | Intercept | -0.09 | 0.31 | 0.22 | 0.03 | 228 | -0.3 | 0.76 | 5 | -1086.06 | 2182.28 | 3.89 | 0.03 |
|  |  |  |  | Potential dispersal rate | -0.01 | 0 |  |  | 228 | -1.55 | 0.12 |  |  |  |  |  |
|  |  |  |  | Local velocity of isotherm shifts | 0.47 | 0.13 |  |  | 160 | 3.53 | 0 |  |  |  |  |  |
| 10 | Local velocity of isotherm shifts | 3rd quartile | - | Intercept | -0.09 | 0.31 | 0.21 | 0.03 | 229 | -0.29 | 0.77 | 4 | -1087.26 | 2182.63 | 4.24 | 0.02 |
|  |  |  |  | Local velocity of isotherm shifts | 0.4 | 0.13 |  |  | 160 | 3.19 | 0 |  |  |  |  |  |
| 11 | Minimum rate | 3rd quartile | Median | Intercept | 0.11 | 0.27 | 0.21 | 0.03 | 229 | 0.4 | 0.69 | 4 | -1087.29 | 2182.68 | 4.29 | 0.02 |
|  |  |  |  | Minimum rate | 0.37 | 0.12 |  |  | 160 | 3.18 | 0 |  |  |  |  |  |
| 12 | Potential dispersal rate (additive) | 3rd quartile | Maximum | Intercept | -0.12 | 0.31 | 0.21 | 0.03 | 228 | -0.4 | 0.69 | 5 | -1086.4 | 2182.96 | 4.57 | 0.02 |
|  |  |  |  | Potential dispersal rate | 0 | 0 |  |  | 228 | -1.31 | 0.19 |  |  |  |  |  |
|  |  |  |  | Local velocity of isotherm shifts | 0.5 | 0.15 |  |  | 160 | 3.42 | 0 |  |  |  |  |  |
| 13 | Potential dispersal rate (interactive) | 3rd quartile | Median | Intercept | -0.14 | 0.31 | 0.22 | 0.04 | 228 | -0.46 | 0.65 | 6 | -1085.79 | 2183.8 | 5.42 | 0.01 |
|  |  |  |  | Potential dispersal rate | 0 | 0.01 |  |  | 228 | -0.01 | 1 |  |  |  |  |  |
|  |  |  |  | Local velocity of isotherm shifts | 0.51 | 0.14 |  |  | 159 | 3.55 | 0 |  |  |  |  |  |
|  |  |  |  | Potential dispersal rate:local velocity of isotherm shifts | 0 | 0 |  |  | 159 | -0.72 | 0.47 |  |  |  |  |  |
| 14 | Potential dispersal rate (interactive) | 3rd quartile | Maximum | Intercept | -0.16 | 0.32 | 0.22 | 0.03 | 228 | -0.5 | 0.61 | 6 | -1086.3 | 2184.82 | 6.44 | 0.01 |
|  |  |  |  | Potential dispersal rate | 0 | 0.01 |  |  | 228 | -0.16 | 0.87 |  |  |  |  |  |
|  |  |  |  | Local velocity of isotherm shifts | 0.52 | 0.15 |  |  | 159 | 3.41 | 0 |  |  |  |  |  |
|  |  |  |  | Potential dispersal rate:local velocity of isotherm shifts | 0 | 0 |  |  | 159 | -0.44 | 0.66 |  |  |  |  |  |
| 15 | Potential dispersal rate | - | Maximum | Intercept | 0.56 | 0.24 | 0.18 | 0 | 228 | 2.36 | 0.02 | 4 | -1092.21 | 2192.52 | 14.13 | 0 |
|  |  |  |  | Potential dispersal rate | 0 | 0 | 0.18 | 0 | 228 | 0.44 | 0.66 |  |  |  |  |  |
| 16 | Potential dispersal rate | - | Median | Intercept | 0.62 | 0.23 | 0.19 | 0 | 228 | 2.68 | 0.01 | 4 | -1092.24 | 2192.57 | 14.19 | 0 |
|  |  |  |  | Potential dispersal rate | 0 | 0 | 0.19 | 0 | 228 | -0.38 | 0.7 |  |  |  |  |  |

| 10 | Potential dispersal rate | - | Max | Intercept | 0.57 | 0.24 | 0.16 | 0 | 259 | 2.33 | 0.02 | 446 | 4 | -1294.46 | 2597.01 | 12.95 |
| --- | --- | --- | --- | --- | --- | --- | --- | --- | --- | --- | --- | --- | --- | --- | --- | --- |
|  |  |  |  | Potential dispersal rate | 0 | 0 |  |  | 259 | 1.1 | 0.27 |  |  |  |  |  |
| 11 | Potential dispersal rate: additive | 3rd quartile | Median | Intercept | 0.29 | 0.29 | 0.15 | 0.01 | 259 | 0.99 | 0.32 | 446 | 5 | -1293.46 | 2597.06 | 13 |
|  |  |  |  | Potential dispersal rate | 0 | 0 |  |  | 259 | -0.63 | 0.53 |  |  |  |  |  |
|  |  |  |  | Local velocity of isotherm shifts | 0.17 | 0.07 |  |  | 184 | 2.33 | 0.02 |  |  |  |  |  |
| 12 | Potential dispersal rate: additive | 3rd quartile | Max | Intercept | 0.25 | 0.29 | 0.15 | 0.01 | 259 | 0.86 | 0.39 | 446 | 5 | -1294.14 | 2598.43 | 14.37 |
|  |  |  |  | Potential dispersal rate | 0 | 0 |  |  | 259 | 0.32 | 0.75 |  |  |  |  |  |
|  |  |  |  | Local velocity of isotherm shifts | 0.15 | 0.08 |  |  | 184 | 2.01 | 0.05 |  |  |  |  |  |
| 13 | Potential dispersal rate: interactive | Mean | Median | Intercept | 0.3 | 0.28 | 0.15 | 0.02 | 259 | 1.07 | 0.29 | 446 | 6 | -1297.59 | 2607.36 | 23.3 |
|  |  |  |  | Potential dispersal rate | 0 | 0.01 |  |  | 259 | 0.04 | 0.97 |  |  |  |  |  |
|  |  |  |  | Local velocity of isotherm shifts | 0.25 | 0.1 |  |  | 183 | 2.62 | 0.01 |  |  |  |  |  |
|  |  |  |  | Potential dispersal rate:velocity of isotherm shift | 0 | 0 |  |  | 183 | -0.46 | 0.64 |  |  |  |  |  |
| 14 | Potential dispersal rate: interactive | 3rd quartile | Median | Intercept | 0.28 | 0.29 | 0.15 | 0.01 | 259 | 0.94 | 0.35 | 446 | 6 | -1298.55 | 2609.3 | 25.24 |
|  |  |  |  | Potential dispersal rate | 0 | 0.01 |  |  | 259 | 0 | 1 |  |  |  |  |  |
|  |  |  |  | Local velocity of isotherm shifts | 0.18 | 0.08 |  |  | 183 | 2.27 | 0.02 |  |  |  |  |  |
|  |  |  |  | Potential dispersal rate:velocity of isotherm shift | 0 | 0 |  |  | 183 | -0.27 | 0.79 |  |  |  |  |  |
| 15 | Potential dispersal rate: interactive | Mean | Max | Intercept | 0.26 | 0.28 | 0.15 | 0.01 | 259 | 0.91 | 0.36 | 446 | 6 | -1299.72 | 2611.62 | 27.56 |
|  |  |  |  | Potential dispersal rate | 0 | 0 |  |  | 259 | 0.22 | 0.82 |  |  |  |  |  |
|  |  |  |  | Local velocity of isotherm shifts | 0.23 | 0.1 |  |  | 183 | 2.17 | 0.03 |  |  |  |  |  |
|  |  |  |  | Potential dispersal rate:velocity of isotherm shift | 0 | 0 |  |  | 183 | -0.2 | 0.84 |  |  |  |  |  |
| 16 | Potential dispersal rate: interactive | 3rd quartile | Max | Intercept | 0.25 | 0.3 | 0.15 | 0.01 | 259 | 0.84 | 0.4 | 446 | 6 | -1300.69 | 2613.56 | 29.5 |
|  |  |  |  | Potential dispersal rate | 0 | 0 |  |  | 259 | 0.12 | 0.9 |  |  |  |  |  |
|  |  |  |  | Local velocity of isotherm shifts | 0.15 | 0.08 |  |  | 183 | 1.83 | 0.07 |  |  |  |  |  |
|  |  |  |  | Potential dispersal rate:velocity of isotherm shift | 0 | 0 |  |  | 183 | 0.06 | 0.96 |  |  |  |  |  |

Note: To assess if results changed when extreme range contractions were included, we refit the 16 candidate models of 5 model types to species’ estimated range shift rates at poleward range edges where local range expansion was expected based on the direction of isotherm velocity, this time excluding extreme negative shifts (n = 391). All models included species as a random effect on the intercept and all variables were continuous measurements in units of km/yr. Models are ranked from lowest to highest AICc value, and significance tests report whether coefficients deviate from 0.

**Table S5.** Summary of model fits across different model types for models fit to local range expansion rates including extreme contractions

| Model type | Lowest model rank | Mean model rank | Cumulative weight |
| --- | --- | --- | --- |
| Potential dispersal rate and local velocity of isotherm shifts (additive) | 1 | 6 | 0.38 |
| Potential dispersal rate and local velocity of isotherm shifts (interactive) | 3 | 8.75 | 0.23 |
| Minimum rate | 4 | 7.25 | 0.26 |
| Local velocity of isotherm shifts | 7 | 8.5 | 0.11 |
| Potential dispersal rate | 15 | 15.5 | 0.00 |

Note: Model ranks and weights based on AIC score of 16 individual models are summarized within each of the five main model types. Models within model type varied in specific parameters used (i.e., 3rd quantile versus mean velocity of isotherm shift, median versus maximum potential dispersal rate; see Table S4 for individual model results).

Dataset S1 (separate file). Raw dispersal distances, dispersal frequencies, and calculated potential dispersal rates.

Dataset S2 (separate file). Information on cause of range contraction for species with extreme contractions at the leading range edge.
